# The major nucleoid-associated protein WHIRLY1 promotes chloroplast development in barley

**DOI:** 10.1101/2024.05.06.592765

**Authors:** Karin Krupinska, Jürgen Eirich, Urska Repnik, Christine Desel, Monireh Saeid Nia, Anke Schäfer, Ulrike Voigt, Bationa Bennewitz, Wolfgang Bilger, Iris Finkemeier, Götz Hensel

## Abstract

WHIRLY1 is a DNA-binding protein of high abundance in chloroplast nucleoids, which have a complex proteome consisting of proteins involved in gene expression and unexpected proteins indicating links to energy production and biosynthetic activities of chloroplasts. In addition, WHIRLY1 has a second localization in the nucleus making it an excellent candidate for chloroplast-to-nucleus communication. To unravel the role of WHIRLY1 for structure and protein composition of nucleoids and its potential involvement in retrograde signaling during chloroplast development, knockout mutants of *HvWHIRLY1* were prepared by site-directed mutagenesis using Cas9 endonuclease. In contrast to mutants of rice and maize, which die after the seedling stage, the barley *why1* mutants survive and produce grains. Leaves of the mutants are initially pale and get green with time (*xantha-to-green* phenotype). However, the chlorophyll content of primary leaves stayed distinctly lower than that of the wild-type leaves, coinciding with a rather heterogeneous plastid population, whereby only 50% developed a rather normal thylakoid membrane system. For comparison, mature foliage leaves had almost normal levels of chlorophyll but a severely reduced photosynthetic capacity.

A proteome analysis of chloroplasts isolated from mature foliage leaves revealed that in the absence of WHIRLY1, the abundances of a considerable fraction of proteins were downregulated. The fraction included multiple nucleoid-associated proteins including components of the transcriptional apparatus. Furthermore, ribosomal proteins, subunits of pyruvate dehydrogenase, CLP protease, ATP synthase, Rubisco and chaperons/chaperonins were found to be downregulated.

In conclusion, the characterization of the barley *why1* mutant plants revealed that WHIRLY1 is not essential for chloroplast development. Rather, it ensures a fast and failure-free progression of chloroplast development by remodeling nucleoids, which serve as assembly platforms for a concerted workflow of the numerous processes required for chloroplast development. Gene expression analyses revealed that the disturbance of chloroplast development is signaled to the nucleus, indicating that WHIRLY1 is not part of the biogenic retrograde signaling of plastids.

## INTRODUCTION

WHIRLY1 belongs to the small family of DNA/RNA-binding proteins called WHIRLIES that have a broad impact on plants’ developmental processes and plant stress resilience (Krupinska et al., 2022; Taylor *et al*., 2022). All genuine WHIRLY proteins have a KGKAAL motif in the WHIRLY domain required for DNA binding (Cappadocia et al. 2012). Most plants, such as barley, have two WHIRLY proteins, whereby WHIRLY1 is targeted to chloroplasts, where it is a major component of the nucleoids (Pfalz et al., 2006; Melonek et al., 2010). WHIRLY1 of barley and other monocot species was shown to induce compaction of nucleoids (Oetke et al., 2022) in line with its proposed function as the eukaryotic architect of chloroplast nucleoids (Kobayashi et al. 2016). For several WHIRLY proteins, multiple localization in two or three DNA-containing compartments has been shown (Krupinska et al., 2022). In barley, WHIRLY1 has been localized in chloroplasts and nuclei within the same cell (Grabowski et al., 2008), making it an attractive candidate mediator of chloroplast-nucleus communication (Krupinska et al. 2020). Protein translocation from chloroplasts to the nucleus has been proposed as one of the multiple modes of retrograde signaling (Bobik and Burch-Smith 2015, Gawronski et al. 2021), which is required for the coordination of gene expression in the nucleus and organelles (Chan et al. 2016, Souza et al. 2017, Pfannschmidt et al. 2020). Biogenic signaling during chloroplast development has been opposed to operational signaling, informing about the functionalities of developed chloroplasts (Pogson et al. 2008). Since *WHIRLY1* knockout mutants of maize (Prikryl et al. 2008) and rice (Qui et al. 2022) are compromised in chloroplast development, WHIRLY1 was designated as a biogenic signal (Chan et al. 2016, Liebers et al. 2022).

Chloroplast nucleoids have a complex protein composition (Melonek et al., 2016). The catalogue of proteins shared by nucleoids and transcriptionally active chromosomes (TAC), which were prepared with different methods from different plant species includes the components of the plastid-encoded RNA polymerase, further proteins involved in DNA-associated processes such as replication, repair, recombination (RRR) besides posttranscriptional processes, in particular translation. In addition to the proteins involved in different steps of gene expression, different nucleoid proteomes share subunits of various metabolic enzyme complexes not expected to reside in nucleoids (Table 2 in Melonek et al. 2016). These unexpected nucleoid-associated proteins (UNAPs) shared by the different nucleoid preparations include 10 ribosomal proteins, four subunits of ATP synthase, four subunits of pyruvate dehydrogenase and acetyl-CoA carboxylase, respectively, besides components required for CO2 fixation, i.e. the large subunit of Rubisco, sedoheptulose biphosphatase, Rubisco activase and chaperonins for the assembly of Rubisco (Melonek et al., 2016).

During chloroplast development, the formation of thylakoids is tightly linked with changes in nucleoid morphology and distribution. While at an early stage of chloroplast development, nucleoids are attached to the inner membrane, where they form a ring-like structure resembling a necklace of pearls; they translocate to thylakoids with advancing chloroplast development (Powikrowska et al., 2014). Nucleoids function in plastid replication and gene expression and were also proposed to serve as platforms for the processing of ribosomal RNA and assembly of the ribosomes (Bohne, 2014).

The attachment of nucleoids at thylakoid membranes was proposed to facilitate the coordination of photosynthesis and gene expression (Powikrowska et al., 2014). In particular, this is important for the remodeling of the photosynthetic machinery by light-dependent changes in the redox state (Pfannschmidt et al., 1999). Likely, the reorganization of the nucleoids during chloroplast development is linked to the establishment of the transcriptional subdomain in the nucleoid that is dominated by the plastid-encoded RNA polymerase proposed to be the bottleneck of chloroplast development (Pfalz and Pfannschmidt, 2013).

The architecture/structure of nucleoids in bacteria is regulated by a group of abundant nucleoid-associated proteins called NAPs, such as HU (heat-unstable protein) and HNS histone-like nucleoid structuring protein), which affect gene expression (Luijsterburg et al., 2006; Dillon and Dorman, 2010). During the evolution of land plants, the NAPs of the cyanobacterial progenitor of plastids were replaced by eukaryotic factors originally involved in the organization of the nuclear chromatin (Sato 2002; Kobayashi et al., 2016). Among these are WHIRLY1 and the MAR-binding-filament-like protein1 (MFP1) proposed to anchor nucleoids to the thylakoid (Meier et al., 1996; Jeong et al., 2003). The compacting impact of barley WHIRLY1 on bacterial nucleoids supports the concept that WHIRLY1 is a major architect of nucleoids (Oetke et al., 2019). Mutants lacking NAPs, such as WHIRLY1, are expected to help to elucidate these proteins’ roles in the expression of plastid genes and photosynthesis (Powikrowska et al., 2014).

When expression of *HvWHIRLY1* was knocked down, the compactness of chloroplast nucleoids was reduced (Krupinska et al., 2014), coinciding with a delay of chloroplast development (Krupinska et al., 2019). *WHIRLY1* transposon insertion mutants of maize show ivory or pale green phenotypes at the seedling stage and do not develop further after the four-leaf stage (Prikryl et al., 2008). Rice mutants with a knockout of *WHIRLY1* have an albino-lethal phenotype and cannot develop after they reach the three-leaf stage (Qiu et al., 2022).

This study addresses the role of WHIRLY1 during chloroplast development. In contrast to the chlorotic and lethal maize and rice *WHIRLY1* knockout mutants (Prikryl et al. 2008, Qui et al. 2022), barley *why1* knockout mutants prepared by site-directed mutagenesis show a xantha-to-green phenotype and survive. Therefore, they are an excellent tool to unravel the actual role of WHIRLY1 during chloroplast development. Dramatic alterations in the proteome of mutant chloroplasts include the group of nucleoid-associated proteins and subunits of supramolecular complexes required for chloroplast development. The results of this study suggest that WHIRLY1 has a structural role, allowing a fast and failure-free progression of chloroplast development by a concerted workflow of the numerous processes required for chloroplast development. The severe disturbances in chloroplast development in the absence of WHIRLY1 are signaled to the nucleus, indicating that WHIRLY1 itself does not function as a biogenic retrograde signal.

## RESULTS

### *why1* KO mutants prepared by Cas9-mediated site-directed mutagenesis

To abolish the function of WHIRLY1 in stable knockout mutants, targeted mutagenesis using the CRISPR/Cas9 technology was employed. Two gRNAs were selected with binding sites before the KGKAAL domain, which is important for DNA binding (Cappadocia et al., 2012). Four independent mutants with pale green leaves at the seedling stage were obtained. All mutations affected the sequence upstream of the WHIRLY domain (Figure 1A, C, Supplemental Figure 1, online). Mutants *why1-2* and *why1-3* have a one nucleotide insertion at position 211, leading to a shift in the reading frame (Fig 1B, D). Mutants *why1*-*1* and *why1*-*4* have deletions of 11 and 25 nucleotides, respectively. All mutants lack the complete WHIRLY1 domain including the KGKAAL motif (Figure 1D, Supplemental Figure 11B, online).

**Figure 1.**
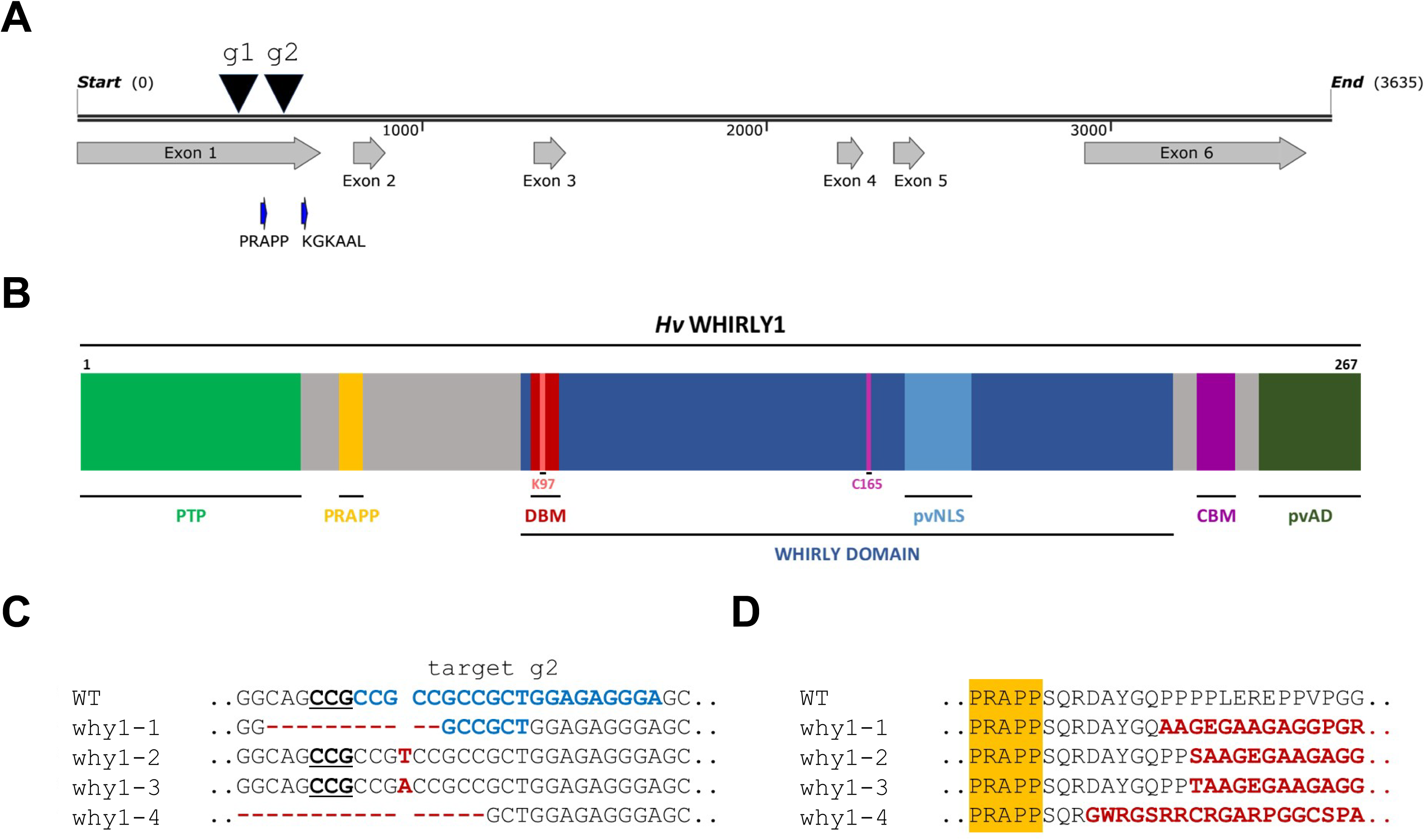
Molecular characterisation of *why1* mutants. **(A)** Genomic organisation of HvWHIRLY1. The two gRNA target sites in exon 1 were indicated by black arrows. In addition, the positions of PRAPP and KGKAAL domains were indicated too. **(B)** Scheme of the WHIRLY1 protein showing different predicted domains (adapted from Krupinska et al., 2022). The WHIRLY domain is marked in dark blue colour. PTP – plastid transit peptide, DBM – DNA-binding motif, pvNLS – putative nuclear localisation signal, CBM – copper-binding motif, pvAD – putative activation domain. **(C)** Sequence comparisons of gRNA2 target site of *why1* mutants with the wild-type sequence. The target site is indicated in blue letters, while PAM is highlighted in bold and underlined. Insertions or deletions (Indels) are shown in red. **(D)** Comparisons of partial wild-type and why1 mutant amino acid sequences at the gRNA2 target site. The PRAPP motif is highlighted in yellow.

A search for potential off-targets of the mutation indicated that the gRNA target sequence was specific for the *WHIRLY1* gene (Supplemental Figure 1A, online). Only the *why1-2* progeny was free of T-DNA among the mutants obtained. For this reason, this mutant line was primarily used in this study. However, the phenotypes of all mutants were similar, and in some experiments, additional mutants were used for comparison.

### Compromised chloroplast development in primary foliage leaves

The leaves of mutant seedlings were pale green at the beginning of leaf development (Figure 2A). The primary foliage leaves of the four mutant seedlings were obviously shorter than those of the wild type (Figure 2A). To investigate whether the kinetics of leaf development is affected by the mutation of *WHIRLY1*, the lengths of the leaves (sheath and blade) were measured on different days after sowing. Mutant leaves were shorter than wild-type leaves at all times of measurement and reached their final length of only 65-75% of the wild-type leaves at the same time as the leaves of the wild type (Fig. 2B).

**Figure 2.**
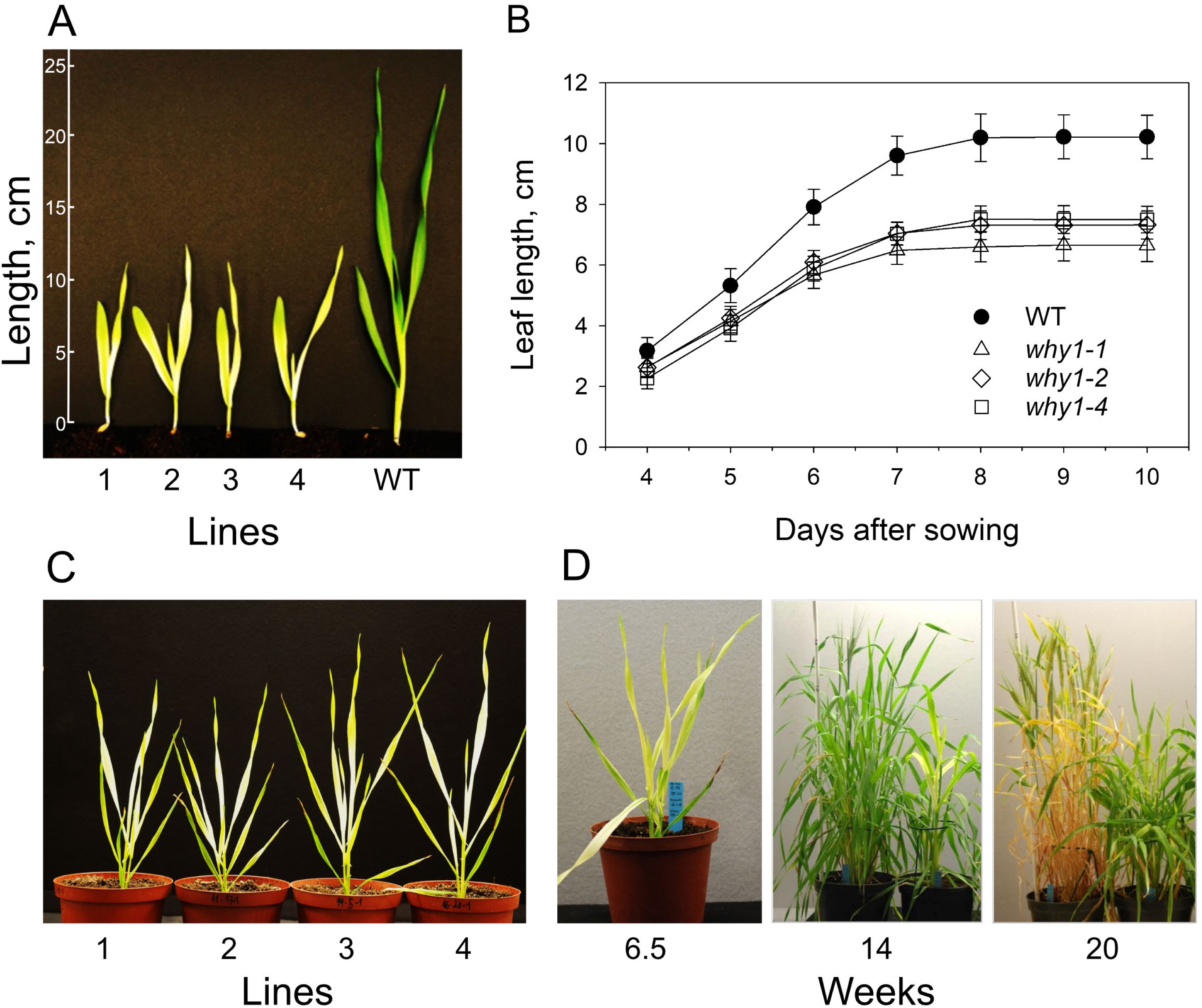
Characterization of leaf development of *why1* mutants. **(A)** Seedlings of four mutants (*why1-1, why1-2, why1-3, why1-4*), compared to a wild-type seedling at 17 days after sowing, **(B)** length of seedlings (cm) measured from the kernel until the tip at different days after sowing, **(C)** mutant plants at 45 das, **(D)** a *why1-4* mutant plant at different weeks after sowing in comparison to a wild-type plant of the same age which after 14 weeks and after 20 weeks is shown at the left, respectively.

Primary foliage leaves of mutant seedlings had light yellow/green colors, whereas the tip of the leaves was green. Foliage leaves showed a similar pigmentation with progressive greening from the tip. According to the nomenclature of barley pigment mutants (von Wettstein, 1959; Nielsen *et al*., 1979; von Wettstein, 2001), the *why1* mutants can be designated as *xantha* mutants that get green with time (Afsson et al. 1938, cited by Rotasperti et al. 2020). This phenotype has been designated *xantha*-to-green. In contrast to the barley *cmf3* mutant (Li et al., 2021), where different parts of the leaf blade simultaneously get green with time, the greening of *why1* leaves begins apparently at the tip of foliage leaves that due to the basipetal growth of cereal leaves contain the eldest cells (Figure 2D). In contrast to foliage leaves, the tip of primary leaves is green from the beginning, indicating that chloroplast development in plastids originating from the embryo is unaffected (Colombo et al., 2008).

HPLC analyses of pigments extracted from segments excised at a position of 1.5 cm below the tip revealed that the mutant leaves had lower chlorophyll content than the wild-type leaves (Figure 3A). Whereas wild-type leaves reached their highest chlorophyll concentration already at eight days after sowing, chlorophyll levels of the mutant leaves continuously increased until 20 days after sowing (*das*) (faster between 6 and 12 *das*), when the chlorophyll content of wild-type leaves already declined due to beginning senescence (Figure 3A).

**Figure 3.**
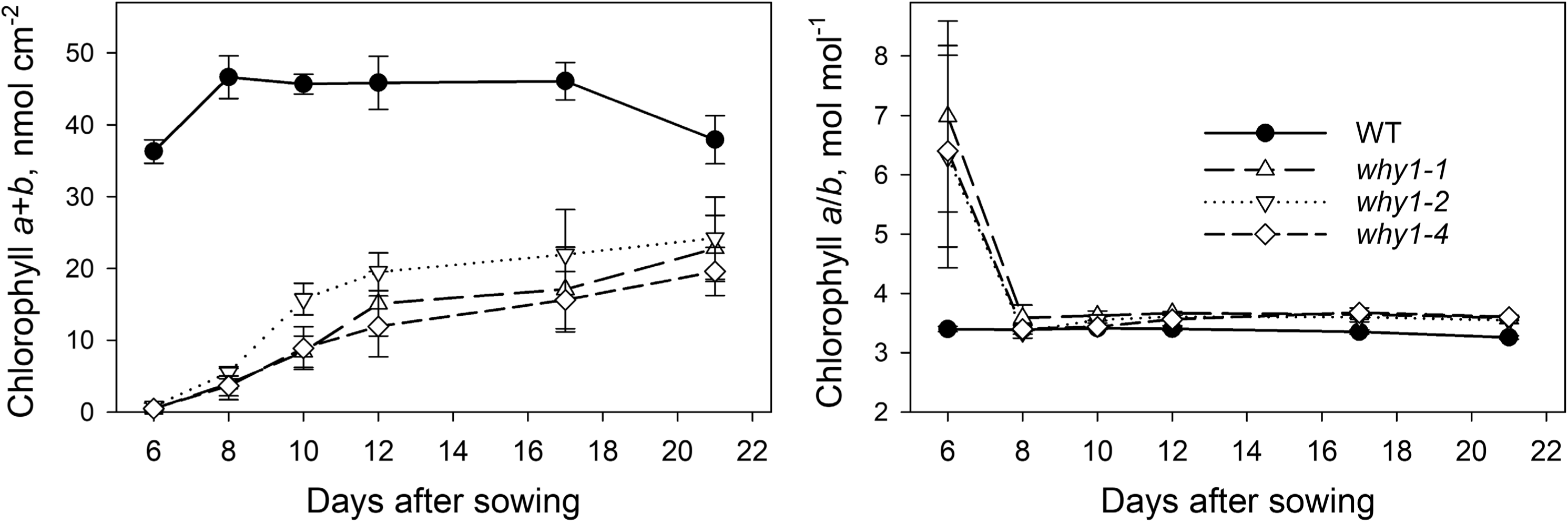
Chloroplast development in primary foliage leaves. **(A)** Chlorophyll content and **(B)** chlorophyll a/b ratio of primary foliage leaves from three mutants (*why1-1, why1-2*, *why1-4)* and the wild type collected in the growth period from day 6 until day 21 after sowing (WT).

Before the senescence-related decline, the chlorophyll content of wild-type leaves (12 das) was more than twice as high as that of the mutant leaves (Figure 3A). At first glance, these results indicate that chloroplast development was simply delayed in mutant leaves. However, even at 21 das, when leaves of all genotypes were fully expanded, and wild-type leaves had already lost 20% of their maximal chlorophyll, the chlorophyll content of mutant leaves was still much lower. Whereas in the primary foliage leaves of the wild type, the maximal chlorophyll content was achieved at the same time when the leaves were fully expanded, the chlorophyll content of mutant leaves continuously increased after the leaves were fully expanded, indicating an uncoupling of leaf development and chloroplast development (compare Figures 2B and 3A).

The high chlorophyll a/b ratio (6-7) determined in mutant leaf extracts at six *das* (Figure 3B) indicates that the plastids were in a very early stage of development, at which the low amount of chlorophyll is primarily used for the formation of reaction center complexes (Akoyunoglou et al., 1966). After 8 days from sowing, the chlorophyll a/b ratio did not differ between mutants and the wild type indicating that at this stage of development the photosynthetic apparatus might have attained a similar composition in all genotypes.

To investigate the stages of the delayed chloroplast development in mutant seedlings in more detail, ultrastructural analyses of cross sections from primary foliage leaves collected at 10 and 17 *das* was performed. Images obtained with the *why1-2* mutant are presented in Figure 3, and images obtained with the *why1-4* mutant are presented in Supplemental Figure 4. At low magnification, it was obvious that mesophyll cells in the sections from leaves collected at 10 das had a dramatically reduced cytoplasm area compared to the large central vacuole (Figure 3C). While the number of plastids was obviously not different between wildtype cells and mutant cells, the size of plastids was much smaller in the mutants compared to the wild type. At higher magnification, it became visible that 10 days after sowing, most mutant plastids contained vesicles and plastoglobules, and only a few contained tiny thylakoid-like structures (Figure 3D). The structural features of most plastids resemble those of plastids in the white stripes of the barley mutant *albostrians*, which is affected by a failure in ribosome biogenesis (Supplemental Figure 4A, online). Intriguingly, different types of plastids were found in one single cell of the *why1* mutants. This feature is in striking contrast to both, cells of the wild type and the *albostrians* mutant, respectively.

Sections obtained from leaves collected at 17 das showed an even more dramatic heterogeneity of plastids. About half of the plastids in one mesophyll cell contained thylakoids (Figure 3C, D). The remaining plastid fraction contained huge vesicles which resembled those observed before in leaves of a transplastomic tobacco mutant lacking the plastid-encoded RNA polymerase (De Santis-Maciossek et al., 1999).

The heterogeneity of the mesophyll plastid population in the *why1* mutant is already visible at 10 das, but it is even more striking at 17 *das* (Supplemental Figure 3, online). It is possible that the vesicles detected in mutant cells at 10 *das* fuse, resulting in the huge vesicles typical for about half of the plastids in mutant cells at 17 *das*. Mutant chloroplasts detected at 17 *das* have no obvious starch granules and a lighter stroma than wild-type ones (Figure 3D).

### Characterization of chloroplasts in green foliage leaves

In contrast to the primary foliage leaves, higher-order foliage leaves of the *why1* mutants attained similar greenness as wild-type leaves (Figure 2D). To investigate whether chloroplast development in mutant leaves is just delayed but ends up at the same stage as in the wild type, chlorophyll content, photosynthesis and chloroplast ultra-structure were analyzed with fully expanded foliage leaves (leaf number 3-4) at the time when they attained their maximal greenness, which was determined by their transmission measured non-invasively with a Dualex device. To attain full greenness, mutant leaves needed a much longer time. To compare chloroplasts from wild-type and mutant leaves, the leaf material had to be collected at different times after sowing. While wild-type leaves have been collected at one month after sowing, mutant leaves have been collected at three months after sowing.

Fully expanded foliage leaves collected three months after sowing from three *why1* mutants almost reached chlorophyll values of wild-type leaves (Figure 5A). Photosynthesis was characterized with the Cas9-free mutant *why1-2* only. While photosystem II efficiency, deter-mined by the chlorophyll fluorescence parameter FV/FM, was almost identical between the leaves of the wild type and *why1-2* mutant, the light-saturated rate of CO2 fixation was strongly reduced (to about 40%) in the mutant leaves compared to wild-type leaves (Figure 4D, E). Whereas light use efficiency at limiting irradiance was not affected in the mutant, A/Ci curves revealed that carboxylation efficiency was reduced by about 60% in the mutant relative to the wild type (Figure 5C-E).

**Figure 4.**
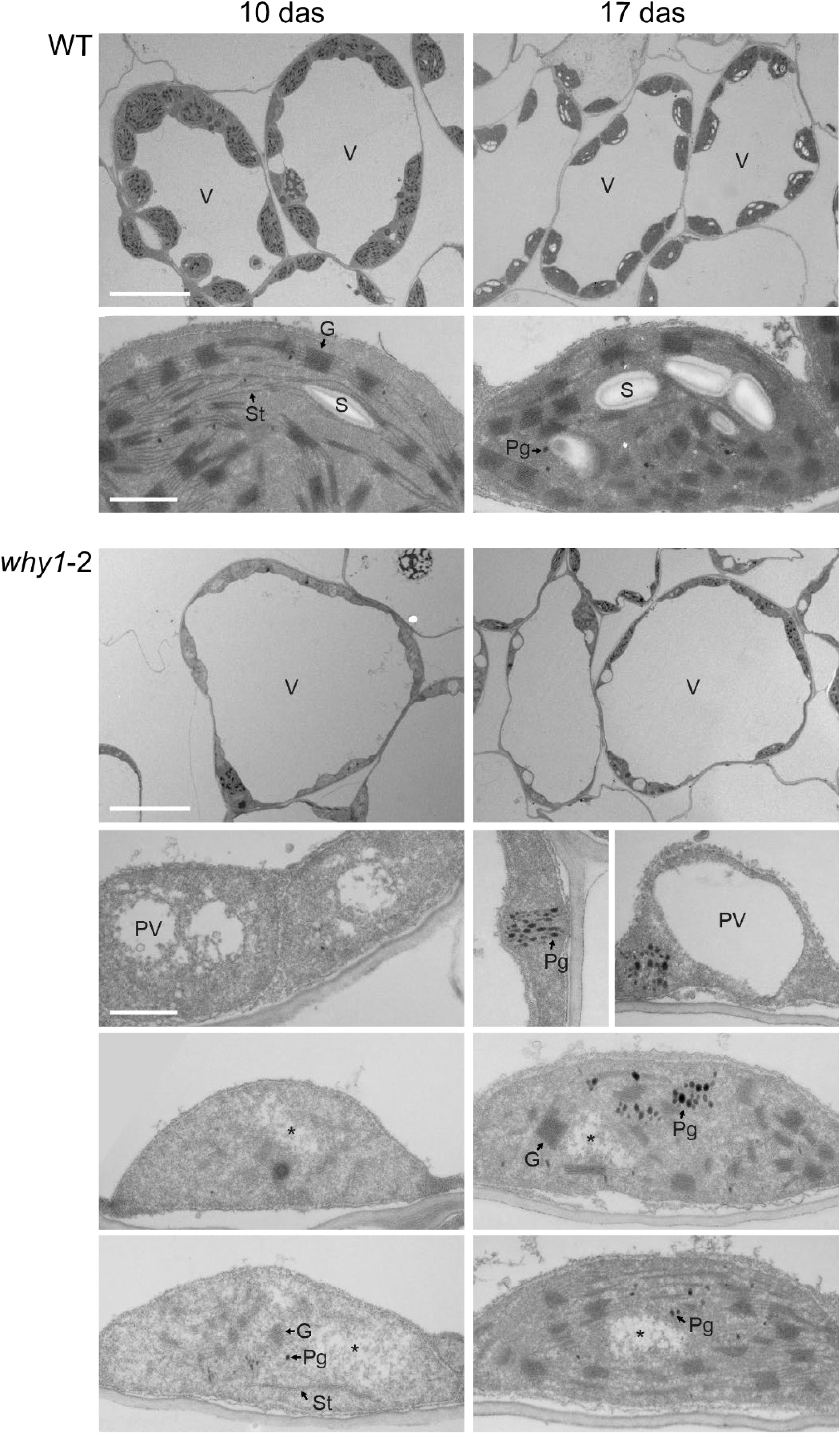
Diversity of chloroplast ultrastructure in WT and *why1*-2 mutant primary foliage leaves at 10 and 17 das. Thin sections of resin embedded leaves were imaged by a transmission electron microscope. G (granum); N (nucleus); Pg (plastoglobule); PV (plastid vesicle); S (starch grain); St (stroma thylakoid); V (central vacuole); asterisk (*) indicates a nucleoid. Scalebar: 10 μm (overviews of cells), 1 μm (chloroplasts).

**Figure 5.**
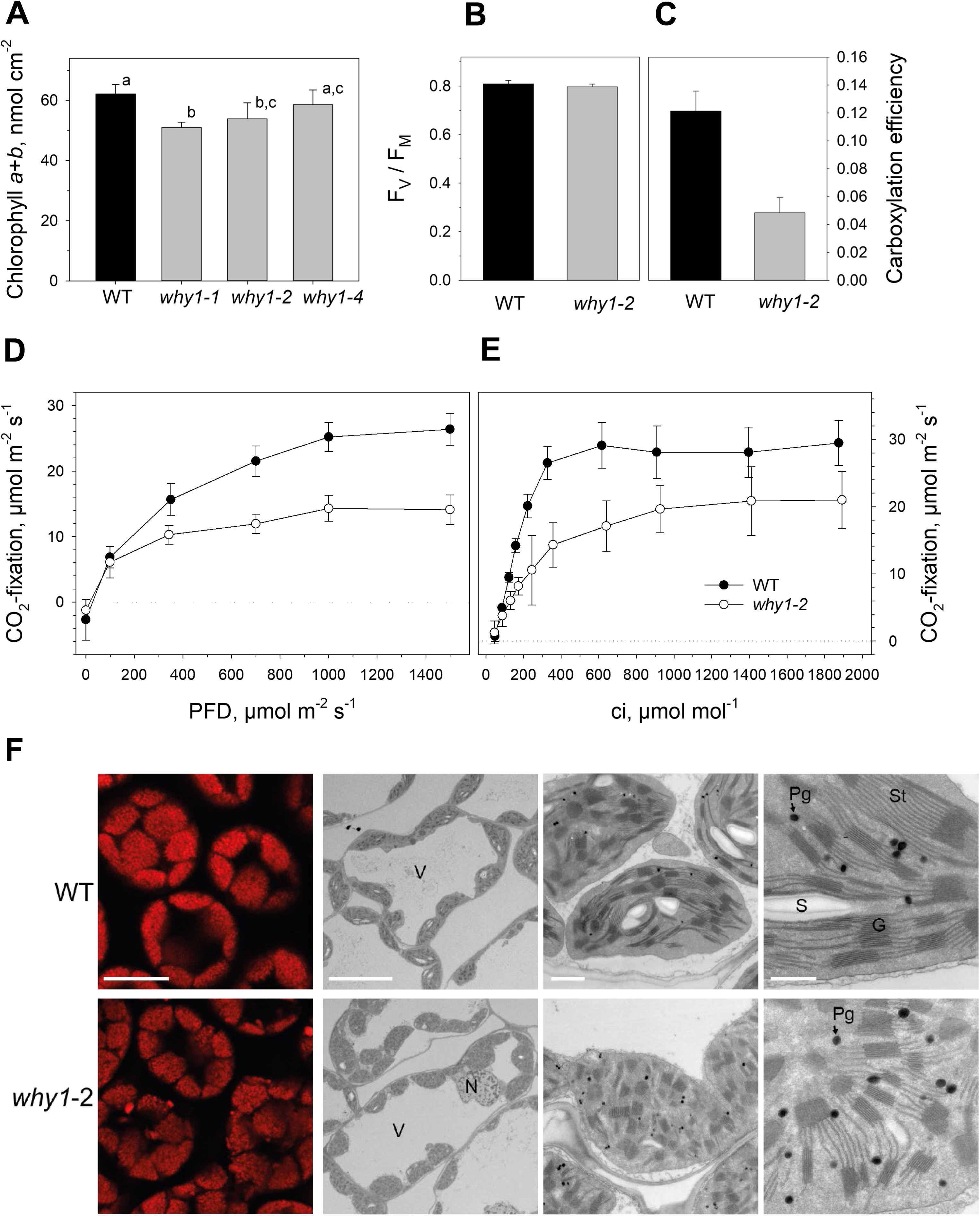
Characterization of fully expanded foliage leaves at the time when they attained maximal greenness. **(A)** Chlorophyll content of leaves collected from the wild-type plants and the three *why1* mutants *(why1-1, why1-2, why1-4*). Pigments have been extracted from segments excised at a position of 2 cm below the tip of foliage leaves. Depicted values are means +/- standard deviations of n=3. **(B, C)** F_V_/F_M_ values (B) and carboxylation efficiencies of foliage leaves collected from the wild type and the mutant *why1-11* **(D, E)** CO_2_ fixation of foliage leaves from the *why1-11* mutant measured at different irradiances (D) and at different CO2 concentrations (E). **(F)** Chloroplast morphology and ultrastructure in WT and why1-2 mutant foliage leaves at 3 months after sowing. Images in the first column represent chlorophyll autofluorescence of chloroplasts in fresh leaves. Images in the other three columns were taken by a transmission electron microscope on thin sections of resin embedded leaves. G (granum); N (nucleus); Pg (plastoglobule); S (starch grain); St (stroma thylakoid); V (central vacuole). Scalebar: 10 μm (fluorescence images, overviews of cells), 1 μm (whole chloroplasts) and 500 nm (details of chloroplasts).

Cross sections cut at 2 cm below the tip of leaves were used for fluorescence microscopy. Fluorescence emission was recorded at 650-720 nm to trace chlorophyll autofluorescence. The images show that the number of chloroplasts was rather similar between mutant and wild-type leaves and that mutant chloroplasts are often associated with tiny fluorescing particles that likely derive from the chloroplasts. Similar chlorophyll-containing particles were found previously under situations of stress and/or senescence and have been interpreted to indicate the degradation of thylakoids (Wang and Blumwald, 2014, Xie et al., 2015). Ultra-structural analyses of cross-sections from the foliage leaves revealed that chloroplasts in the mutant mesophyll cells contain more plastoglobules than the wild-type chloroplasts. This feature might indicate that thylakoids were remodeled or degraded (Rottet et al., 2015). In line with the reduced photosynthetic activity, only a few chloroplasts contained starch granules (Figure 4G, Supplemental Figure 6, online). Like the chloroplasts of primary foliage leaves at 17 *das*, mutant chloroplasts of foliage leaves had a lighter stroma than in the wild-type chloroplasts (Figure 4G). Images show that grana stacks of the mutant chloroplasts are not as uniform as in the wild type and have reduced regularity in their arrangement (Figure 5G).

### Nucleoid morphology and plastid DNA content of primary leaves and green foliage leaves

WHIRLY1 is a major component of chloroplast nucleoids (Pfalz et al. 2006, Melonek et al. 2010). Staining of DNA of chloroplasts in the barley *WHIRLY1* knockdown line W1-7, having only a minute amount of WHIRLY1 revealed that the protein plays a role in the compaction of nucleoids (Krupinska et al., 2014) as confirmed by overexpression of *HvWHIRLY1* in *E. coli* (Oetke et al. 2022). To investigate the morphology of nucleoids in the *why1* knockout mutants, cross-sections from primary foliage leaves collected at 10 *das* and from mature foliage leaves were stained with SYBR Green, respectively. In primary foliage leaves of the mutants, the fluorescence emission of chlorophyll was much weaker than in leaves of the wild type. Images show that the green fluorescence signals from the stained nucleoids were larger in the mutant chloroplasts than in wild-type chloroplasts (Figure 5A). Nucleoid signals in the primary leaves of the mutants were not as heterogeneous as in the chloroplasts of the barley knockdown line W1-7, which still contained a few tiny compact nucleoids (Krupinska et al., 2014). Compared to the primary foliage leaves of mutants, fluorescence signals from mutant chloroplast nucleoids in foliage leaves were smaller (Figure 5B, C). Nevertheless, in both primary foliage leaves and mature foliage leaves of mutant plastids, the areas of SYBR Green signals were larger than in wild-type chloroplasts (Figure 5C).

For comparison, the ratio between plastid DNA content and nuclear DNA was quantitatively determined by real-time PCR employing specific primers for two plastid genes, i.e. *petD* and *psbA* and the nuclear gene *RBCS,* respectively. In primary foliage leaves of the mutant why1-2, the relative plastid DNA content was more than twice as high as in wild-type leaves (Figure 6A). The same result was obtained with the knockdown line W1-7 containing only about 1% of WHIRLY1 protein (Krupinska et al., 2014). In foliage leaves from the *why1* mutant, the relative plastid DNA content was even up to five-fold higher than in comparable wild-type leaves (Figure 6B).

**Figure 6.**
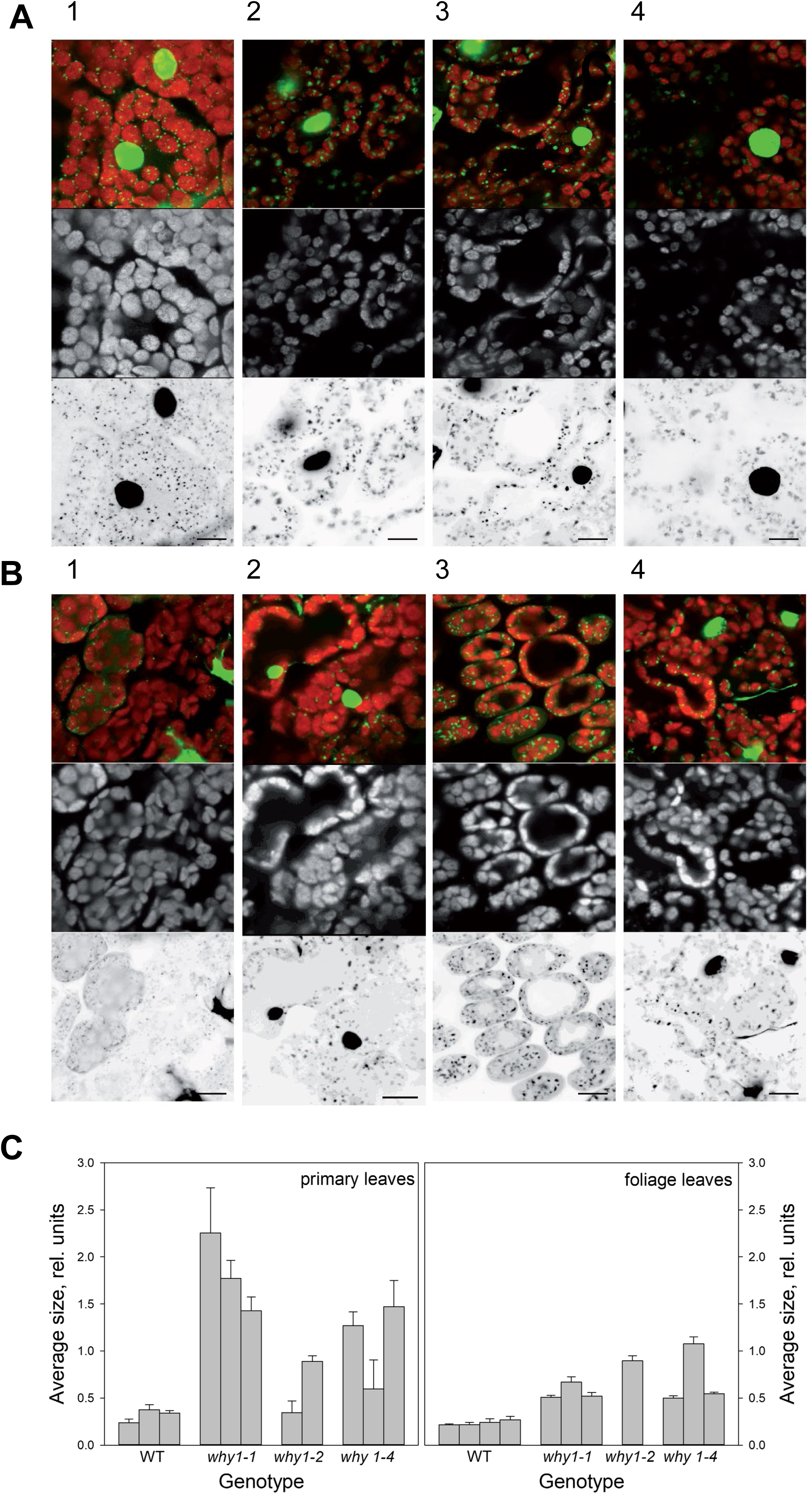
Morphology of plastid nucleoids in the wild type (1, 5) and three *why1* mutants: *why1-1* (2, 6*), why1-3* (3, 7) and *why1*-*4* (4, 8). Leaf sections have been stained with SYBR Green. **(A)** Panels 1-4 show images of sections from the middle of primary foliage leaves, **(B)** panels 5-8 show images from sections prepared from mature foliage leaves cut at positions of 2 cm below the tip. On the top of A and B, signals of the green and red channels were merged. In the middle, chlorophyll fluorescence is shown in light grey and at the bottom, green signals of DNA are shown in black. Excitation was done by an argon laser line 488 (5% power). Emission was detected between 510-570 nm (HV750) and 690-760 nm (HV480). Bars represent 10 µM.

### Levels of RNAs

Chloroplast development is associated with specific changes in plastid DNA transcription, which can be attributed to changes in the ratio of the two plastid localized RNA polymerases (Börner et al., 2015). While the nuclear-encoded RNA polymerase (NEP) is dominant in proplastids and transcribes genes encoding the subunits of the plastid RNA polymerase (PEP), the more efficient plastid-encoded RNA polymerase gains increasing importance during chloroplast development. In barley primary foliage leaves, the development-related changes in transcription have been shown by run-on transcription assays and RNA accumulation in segments derived from different parts of the basipetal leaf (Baumgartner et al., 1989, Rapp et al., 1992). To investigate whether the disturbance in chloroplast development of *why1* mutants is caused by alterations in plastid gene expression, mRNA levels of selected plastid genes were determined by quantitative real-time RT-PCR. Among the genes selected are some that are thought to be primarily transcribed by NEP and some that are thought to be primarily transcribed by PEP.

Levels of ribosomal RNAs, which mainly depend on PEP transcription (Demarsy et al., 2012, Börner et al., 2015), were reduced by 40% in the case of 16S rRNA and even by about 60% in the case of 23S rRNA (Figure 8A). The results are similar between primary foliage leaves on one hand and mature foliage leaves collected at different times after sowing on the other hand. One of the few barley plastid genes transcribed exclusively by PEP is *psbE* (Zhelyazkova et al., 2012). While its mRNA level is not altered in the absence of WHIRLY1, the mRNA levels of genes transcribed by NEP, i.e. *rpoB*, *rpoA* and *clpP*, are highly elevated in the mutant compared to the wild type (Figure 8B). Also, mRNA levels of *rpl2* and *atpF* that are likely transcribed (having transcription start sites for both polymerases) by both polymerases were higher in the mutant leaves. The highest level has been observed for the NEP derived *rpoB* mRNA at the early stage of primary leaf development, i.e. at 10 *das* (Figure 8B). To elucidate whether transcription of plastid genes by PEP is specifically affected in the absence of WHIRLY1, run-on transcripts were hybridized to 23 specific plastid genes dotted in different concentrations on a nylon filter as described earlier (Falk et al., 1993, Melonek et al., 2010). Among the genes were *psbE* and *ndhH,* which in barley have promoters containing only transcription starts sites for PEP (Zhelyazkova et al., 2012). The hybridization signal intensities of these transcripts compared to the signal intensities of *rbcL* transcripts and *rpoB/C1* transcripts which are synthesized from NEP transcription start sites (Zhelyazkova et al., 2012) were barely altered (Supplemental Figure 5). Opposite to the ratio of 16SrRNA and 23SrRNA levels in the mutant (Figure 8A), the relative transcription level of 16SrDNA was reduced compared to 23SrDNA transcription (Supplemental Figure 5). Taken together the results of the run-on transcription assays indicated that the changes in transcript abundance are not caused by specific changes in the transcriptional activities. An estimation of the hybridization signal intensities obtained with run-on transcripts from wild-type and mutant plastids (Supplemental Figure 5) indicated that the overall plastid transcriptional activity in the mutant is only 20% of that of the wild type (Table in Supplemental Figure 5).

The differences in mRNA levels of the *why1* mutant align with those detected in primary foliage leaves of the WHIRLY1 knockdown line W1-7 (Krupinska et al. 2019). In both lines, the ratio of NEP-derived transcripts to PEP-derived transcripts was higher than in the wild type. Taken together, the results are consistent with the role of WHIRLY1 in the acceleration of chloroplast development. In the barley *WHIRLY1* knockdown line W1-7, the high relative level of plastid DNA coincided with high expression of the gene encoding plant organelle DNA polymerase (POP) (Krupinska et al., 2014). Besides, in the primary foliage leaves of the *why1* mutant, the expression of *POP* was higher in the mutant than in the wild type. While the level was 12-fold higher in 10 *das* leaves, it was only about 7-8-fold higher in 17 *das* leaves. Moreover, the expression of *RPOT* encoding NEP was enhanced in mutant leaves of different developmental stages (Figure 8C). In addition, the mRNA levels of selected nuclear genes encoding further nucleoid-associated proteins with a second localization in the nucleus (Krupinska et al. 2020) were determined: MFP1, SVR4 (RBC, MRL7), SVR4-like (NCP, MRL7-L) and PTAC12, which is a protein associated with PEP and hence named also PAP5 (Pfannschmidt, 2015). Moreover, PTAC12 is identical to HEMERA, playing a role in phytochrome-dependent light signalling (Nevarez et al., 2017). Whereas MFP1 had a high expression level (12-fold and 7-8-fold) in primary foliage leaves, its expression was not detectable in mature foliage leaves. The expression levels of *SVR4* and *SVR4-like* were slightly enhanced (up to 5-fold) in the mutant’s primary foliage leaves and foliage leaves.

Compared to the wild type, the expression level of *PTAC12* was 11-fold higher in the primary foliage leaves of the mutant at 10 *das* and about 7-fold higher in the primary foliage leaves at 17 *das*. Whereas the *PTAC12* expression level was enhanced to the same extent in mature foliage leaves collected at 49 das, its expression declined in the elder foliage leaves. For all nuclear genes tested, the increase in the level depended on the age of the leaves, whereby the relative levels were highest in the young primary foliage leaves.

### The proteome of mutant chloroplasts is deficient in numerous nucleoid-associated proteins

Chloroplasts were prepared in four biological replicates and processed in parallel from mature foliage leaves of the wild type and the two mutants, *why1-2* and *why1-4*, respectively. Proteins were extracted from chloroplasts and digested into peptides using trypsin. Equal amounts of purified peptides were analysed by quantitative label-free high resolution mass spectrometry. In total, 2049 chloroplast protein groups were detected in the dataset, with 2041 in chloroplasts of both the *why1-2* mutant and the *why1-4* mutant (Figure 8A, B, Supplemental Table 2, online). While in the *why1-2* chloroplast proteome, 258 proteins were up-regulated, 708 proteins were down-regulated compared to WT, in the *why1-4* mutant 307 chloroplast proteins were up-regulated, and 764 proteins were down-regulated. The correlation of 0.85 between the two mutant proteomes (Figure 8C) indicates that the effect of the mutation was slightly different in the two lines. While four and five peptides of WHIRLY1 (in 3 and 1 replicates, respectively) were detected in the wild-type samples, no peptides of the POI were found in any of the mutant lines. Both proteomes were screened for several functional groups of chloroplast proteins.

Considering that WHIRLY1 is a major architect of nucleoids, its loss was expected to have an impact on the protein composition of nucleoids, which share many proteins with the so-called transcriptionally active chromosome (TAC) (Pfalz et al. 2006, Melonek et al. 2016). The proteome analysis revealed that the loss of WHIRLY1 resulted in dramatic changes in the protein repertoire of nucleoids: while most nucleoid-associated proteins (NAPs) were found to be down-regulated, only a few NAPs were up-regulated (Figure 8D). Regarding the delay in chloroplast development of the mutants and its compromised photosynthesis, the abundance of proteins involved in chlorophyll biosynthesis and photosynthesis was of interest. The box plot shows that all members of the chlorophyll biosynthesis group were downregulated, while most photosynthesis-associated proteins were up-regulated in the mutant proteomes (Figure 8D).

To identify NAPs affected by the loss of WHIRLY1, the Arabidopsis gene identifiers of the catalogue presented in Melonek et al. (2016) were used to screen the list of homologous proteins detected in barley chloroplasts (Supplemental Data, Table X). This catalogue contains 96 proteins shared by several proteomes published for different preparations of nucleoids and transcriptionally active chromosomes from different plants (Melonek et al. 2016). As expected, the catalogue contains factors involved in processes associated with DNA, i.e. replication, repair, recombination, and transcription. Numerous proteins in this group have at least two names because they had been identified in transcriptionally active chromosomes (TAC) and were also found to be associated with PEP (PAPs). Furthermore, this catalogue contains numerous proteins detected consistently in nucleoid and TAC preparations but do not have apparent functions associated with DNA. These unexpected nucleoid-associated proteins (UNAPs) comprise subunits of large complexes such as the ribosome, ATP synthase, pyruvate dehydrogenase, and the CLP protease complex. Moreover, enzymes of the Calvin cycle and the translation elongation factor EF-TU are in this group of proteins (Melonek *et al*., 2016). Further subunits of these complexes were detected in the group of down-regulated proteins of the mutant chloroplast proteome (Table 1). In the context of CO2 fixation, besides the large subunit of Rubisco, the small subunit and five further enzymes of the Calvon-Bassham-Benson (CBB) cycle were found to be down-regulated (Table 1). Interestingly, RAF1, a factor required to assemble Rubisco (Xia et al., 2020) and CP12, was strongly downregulated. CP12 has been described as a protein linker for supramolecular complex assembly (Graciet et al., 2003). The ATP synthase subunit CLB160 was first described as an assembly factor of CF0 and has recently been shown to be required to assemble CF0 and CF1 (Reiter et al., 2023). Besides the chaperone CLPC1, 12 subunits of the CLP protease complex were detected in the chloroplast proteome and were downregulated in the mutant. In addition to the 10 ribosomal proteins listed in the NAP catalogue and found to be down-regulated in the mutant, only one additional ribosomal protein, Rpl2-A, was detected in the chloroplast proteomes. It is downregulated in the mutant as well. Besides the ribosomal proteins, numerous factors involved in ribogenesis (Schmid and Meurer 2023) were down-regulated (Supplemental Table 3, online). Furthermore, several subunits of the plastid chaperonin system, i.e. CPN60, CPN20 and CPN10 (Trösch et al. 2015) are downregulated in the mutant chloroplast proteomes (Table 1). In an earlier study with maize, chaperonins CPN60A/B, CLPC1 have been both enriched in the WHIRLY1 immunoprecipitates from nucleoids (Majeran et al., 2012, Supplemental Table 6) whose composition otherwise hardly differed from total nucleoids (Majeran et al., 2012).

**Table 1.** Relative abundances in the mutant chloroplast proteomes of subunits belonging to multisubunit complexes containing unexpected nucleoid-associated proteins (UNAPs) (Melonek et al. 2016). Rows listing UNAPs of the NAP catalogue (Melonek et al.2016) are highlighted in grey. The relative abundances in the two chloroplast proteomes are indicated by log_2_ fold changes. Furthermore, the occurrence of immunoprecipitates obtained from nucleoids with the maize WHIRLY1 antibody (Supplement of Majeran et al. 2012) are indicated. The highest abundance values in co-immunoprecipitates deduced from the Supplemental Table S6 of Majeran et al. (2012): *NadjSPC x 1000 are indicated in the right column. PDH=pyruvate dehydrogenase, CLP=CLP plastid protease, CBB=Calvin Bassham Benson cycle, n.d.=not detected.

| complex,<br>function | subunit | gene number<br>Arabidopsis | log<sub>2</sub> FC why1-2 | log<sub>2</sub> FC<br>why1-4 WT | co-IP (α-ZmW1)* |
| --- | --- | --- | --- | --- | --- |
| **translation** |  |  |  |  |  |
| ribosome | RPL1 | AT3G63490 | -0.8 | -1.0 | - |
|  | Rpl2-A | ATCG00830 | -6.3 | -1.1 | 8.5 |
|  | Rpl2-B | ATCG01310 | -2.5 | -0.7 | 8.0 |
|  | RPL3 | AT2G43030 | -1.1 | -0.9 | 8.0 |
|  | RPL4 | AT1G07320 | -1.4 | -2.1 | 16.6 |
|  | RPL29 | AT5G65220 | -2.4 | -2.1 | 0.3 |
|  | RPL12-C | AT3G27850 | -2.3 | -1.0 | 0.4 |
|  | Rps2 | ATCG00160 | -2.1 | -3.9 | 3.9 |
|  | Rps3 | ATCG00800 | -1.6 | -1.8 | 5.5 |
|  | Rps4 | ATCG00380 | -5.2 | -4.4 | 9.8 |
|  | RPS5 | AT2G33800 | -3.0 | -2.5 | 2.6 |
| elongation | EF-TU, TUFA | AT4G20360 | -1.8 | -2.6 | 5.4 |
| **CO<sub>2</sub> fixation** |  |  |  |  |  |
| Rubisco | RbcL | ATCG00490 | -8.6 | -9.5 | 0.7 |
| Rubisco | RBCS | AT1G67090 | -5.4 | -5.1 | n.d. |
| Rubisco (R) | RBCS3B | AT5G38410 | -3.9 | -4.3 | n.d. |
| R. activase | RCA | AT2G39730 | -2.1 | -2.3 | 0.9 |
| R. assembly | RAF1 | AT3G04550 | -5.0 | -6.3 | - |
| protein linker | CP12 | AT3G62410 | -5.7 | -3.1 | - |
| CBB enzyme | SBPase | AT3G55800 | -2.8 | -3.6 | - |
| CBB enzyme | GAPA | AT1G12900 | -2.5 | -2.6 | 3.2 |
| CBB enzyme | GAPB | AT1G42970 | -2.0 | -2.6 | 5.1 |
| CBB enzyme | Transketolase | AT3G60750 | -2.4 | -3.4 | n.d. |
| CBB enzyme | PRK | AT1G32060 | -1.9 | -2.6 | n.d. |
| CBB enzyme | CFBP1 | AT3G54050 | -3.0 | -2.9 | n.d. |
| **ATP-synthase** |  |  |  |  |  |
| CF1 | AtpA | ATCG00120 | -4.1 | +0.9 | 4.9 |
| CF1 | AtpB | ATCG00480 | -9.4 | -1.7 | 2.5 |
| CF1 | AtpE | ATCG00470 | -0.3 | -0.2 | n.d. |
| CF<sub>0</sub> chaperon | CGL160 | AT2G31040 | -1.3 | -0.2 | n.d. |
| **CLP protease** |  |  |  |  |  |
| chaperon | CLPC1 | AT5G50920 | -8.2 | -6.9 | 7.3 |
|  | CLPP4 | AT5G45390 | -8.5 | -5.3 |  |
|  | CLPP3 | AT1G66670 | -5.9 | -6.3 |  |
|  | CLPP5 | AT1G02560 | -5.8 | -4.1 |  |
|  | CLPP6 | AT1G11750 | -5.8 | -9.7 |  |
|  | CPLP (pt) | ATCG00670 | -5.8 | -5.4 |  |
|  | CLPR1 | AT1G49970 | -5.7 | -5.3 | 0.3 |
|  | CLPR2 | AT1G12410 | -4.9 | -4.3 | n.d. |
|  | CLPR4 | AT4G17040 | -3.6 | -6.5 | n.d. |
|  | CLPB | At5G15450 | -5.2 | -4.3 | 0.3 |
|  | CLPD | AT5G51070 | -1.6 | -3.0 | n.d. |
|  | CLPR3 | AT1G09130 | -5.9 | -2.6 | n.d. |
| PDH | E1-BETA-1 | AT1G11090 | -5.4 | -3.8 | n.d. |
|  | E1-ALPHA-1 | AT1G01090 | -5.0 | -2.8 | n.d. |
|  | LTA2 (LTH2) | AT3G25860 | -3.9 | -4.7 | n.d. |
| **chaperonin** |  |  |  |  |  |
|  | CPN60A | AT2G28000 | -6.2 | -9.0 | 1.3 |
|  | CPN60B | AT3G13470 | -4.2 | -4.1 | 1.1 |
|  | CPN20 | AT5G20720.1 | -8.2 | -7.0 | n.d. |
|  | CPN10 | AT3G60210 | -7.3 | -10.1 | n.d. |

## DISCUSSION

The albino or ivory phenotypes of the lethal *why1* mutants of maize and rice indicate that WHIRLY1 plays a fundamental role in chloroplast and overall plant development (Prikryl et al., 2008; Qiu et al., 2022). Regarding the high sequence similarities and conserved motif compositions among the monocot WHIRLY1 sequences (Krupinska et al., 2022), it has been expected that the barley *WHIRLY1* knockout mutantphenotype is similar to that of the two other monocot *why1* mutants. However, the phenotype of the *Hvwhy1* mutant is more complex. In contrast to the *why1* mutants of rice (Qie et al. 2022) and maize (Prikryl et al., 2008), the barley *why1* mutants survive and produce grains. It is likely that in barley the loss of WHIRLY1 is compensated by one or several other proteins having partly overlapping functions with WHIRLY1 during chloroplast development.

### Chloroplast development is compromised

The slow increase in chlorophyll content of primary foliage leaves suggests that chloroplast development is delayed. Ultrastructural analyses, however, revealed that chloroplast development is variegated at the subcellular level. During normal chloroplast development in the light, vesicles from the inner envelope fuse to build up the thylakoid membrane system (von Wettstein, 2001). Thereby, all plastids in one cell are at the same stage of development. In contrast, in sections from barley *why1* mutant primary foliage leaves collected at 10 *das*, the same cell contained plastids of different developmental stages. Besides plastids with vesicles and plastoglobules, some plastids were observed to contain tiny thylakoids. The heterogeneity of the plastids is even more pronounced in mesophyll cells of leaves collected one week later. Plastids with huge vesicles were found here besides chloroplasts with a rather normal thylakoid membrane system. It is evident that in the absence of WHIRLY1, the development of the numerous plastids in one cell lacks synchronization.

The huge vesicles might be formed by membrane lipids that do not form bilayers without the appropriate proteins required for membrane organization. Thylakoids contain a large proportion of polyunsaturated galactolipids known as non-bilayer lipids (Vothknecht and Westhoff, 2001). It is likely that the huge vesicles in plastids are produced when membrane lipid biosynthesis proceeds while the abundance of the proteins required for membrane stacking (Vothknecht and Westhoff, 2001) is too low, either due to an impaired protein synthesis capacity of the plastids and/or reduced import of proteins from the cytosol. Plastids with huge vesicles have also been observed in the leaves of the albino tobacco transplastomic mutants lacking the plastid-encoded RNA polymerase (PEP) Field (De Santis-Maciossek et al., 1999). Due to the lack of PEP, plastid protein synthesis is reduced and the protein deficit prevents thylakoid stacking.

### Plastid transcript levels are enhanced in barley *why1* mutants

Plastids have two different types of RNA polymerases. Besides the plastid-encoded RNA polymerase (PEP) of procaryotic origin, they have a nucleus-encoded RNA polymerase resembling the polymerases of phages (NEP) (Liere et al., 2011). The polymerases use different promoters. Analyses of transcripts showed that both polymerases are present at all stages of development in all plant tissues (Börner et al., 2015). Chloroplast development depends mainly on transcription by PEP which is more efficient than NEP and provides the high amounts of transcripts required for a fast build-up of the photosynthetic apparatus (Liere et al., 2011) whose core subunits are encoded by plastid genes, i.e. *psbA* and *psaA/B*. The biosynthesis of PEP itself depends on the activity of NEP. In barley, the *rpoB* gene encoding the ß-subunit of PEP is the only plastid gene that NEP exclusively transcribes (Zhelyazkova et al., 2012; Börner et al., 2015). The level of *rpoB* transcripts, as well as the levels of other transcripts produced by the involvement of NEP, are enhanced in the barley *why1* mutant leaves (Figure 7). The imbalance in transcript levels could be caused by reduced activity of PEP. However, run-on transcription assays with isolated plastids revealed that both RNA polymerases work with about the same efficiency which is reduced in the mutant plastids (signal intensities are lower although the same number of plastids has been used).

**Figure 7.**
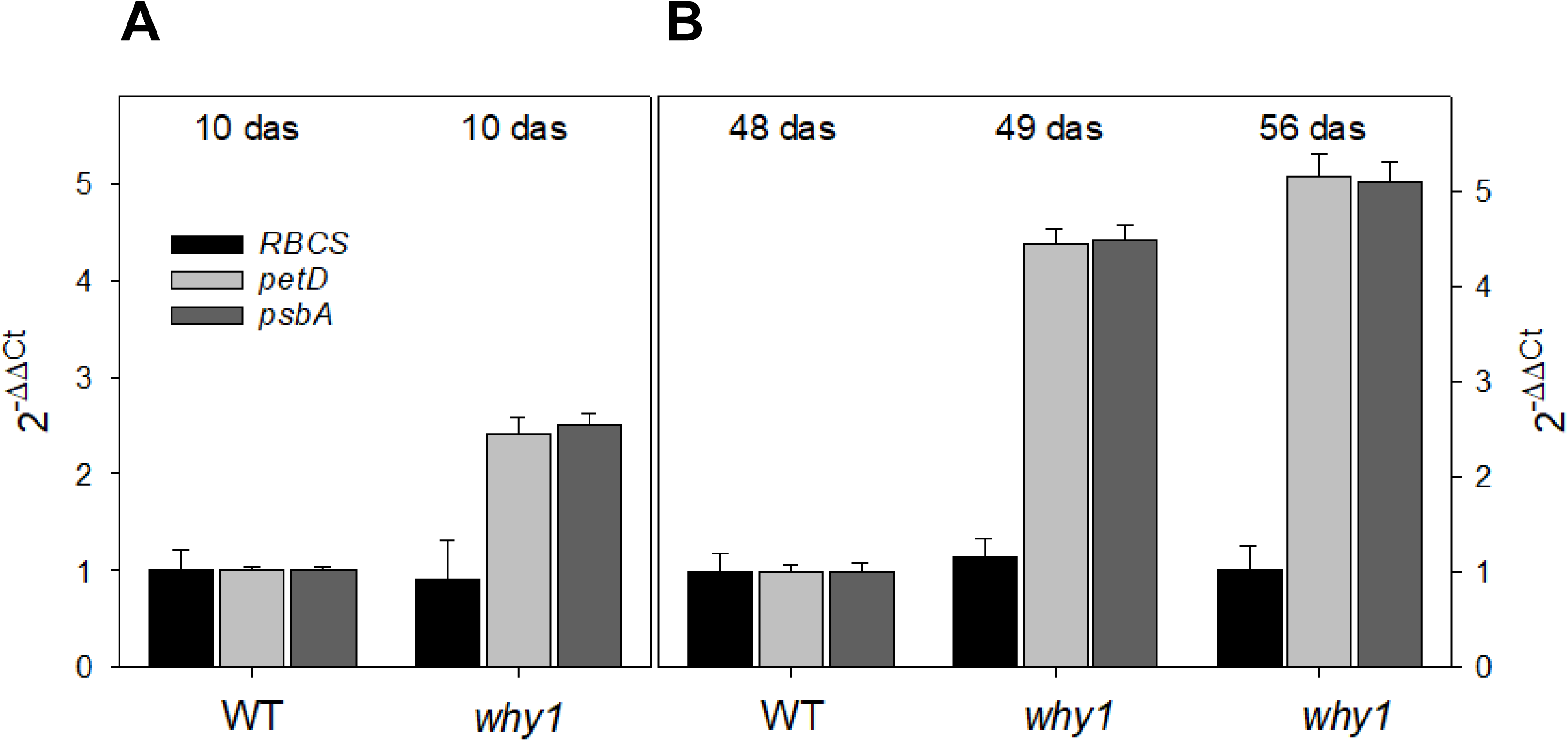
Relative plastid DNA (ptDNA) content in comparison to nuclear DNA (nDNA) in leaves of the wild type and the *why1-2 (why1)* mutant. The relative abundance of ptDNA was determined by qPCR using primers specific for the plastid genes *petD* and *psbA*. Nuclear DNA was amplified with primers specific to the *RBCS* gene. **(A)** levels of ptDNA and nDNA in primary foliage leaves of wild type and the *why1-4* mutant at 10 das, **(B)** levels of ptDNA and nDNA in green foliage leaves collected at different times after sowing of the WT (48 das) and the *why1*-*2* mutant (49, 56 and 66 das). DNA levels were normalized to the level of 18S DNA. Data for the primary foliage leaves of the wild type were set to 1 and data for mutants were calculated relative to the wild type.

While the plastid-encoded PEP core subunits and nuclear-encoded proteins associated with PEP (PAPs) have low abundances in the proteome of the mutant chloroplasts (Figure 9), NEP was not detectable. It is indeed well-known that NEP has a low abundance (Liere et al., 2011). It is likely, that WHIRLY1 is required for assembly of the complete transcriptional apparatus in plastids.

**Figure 8.**
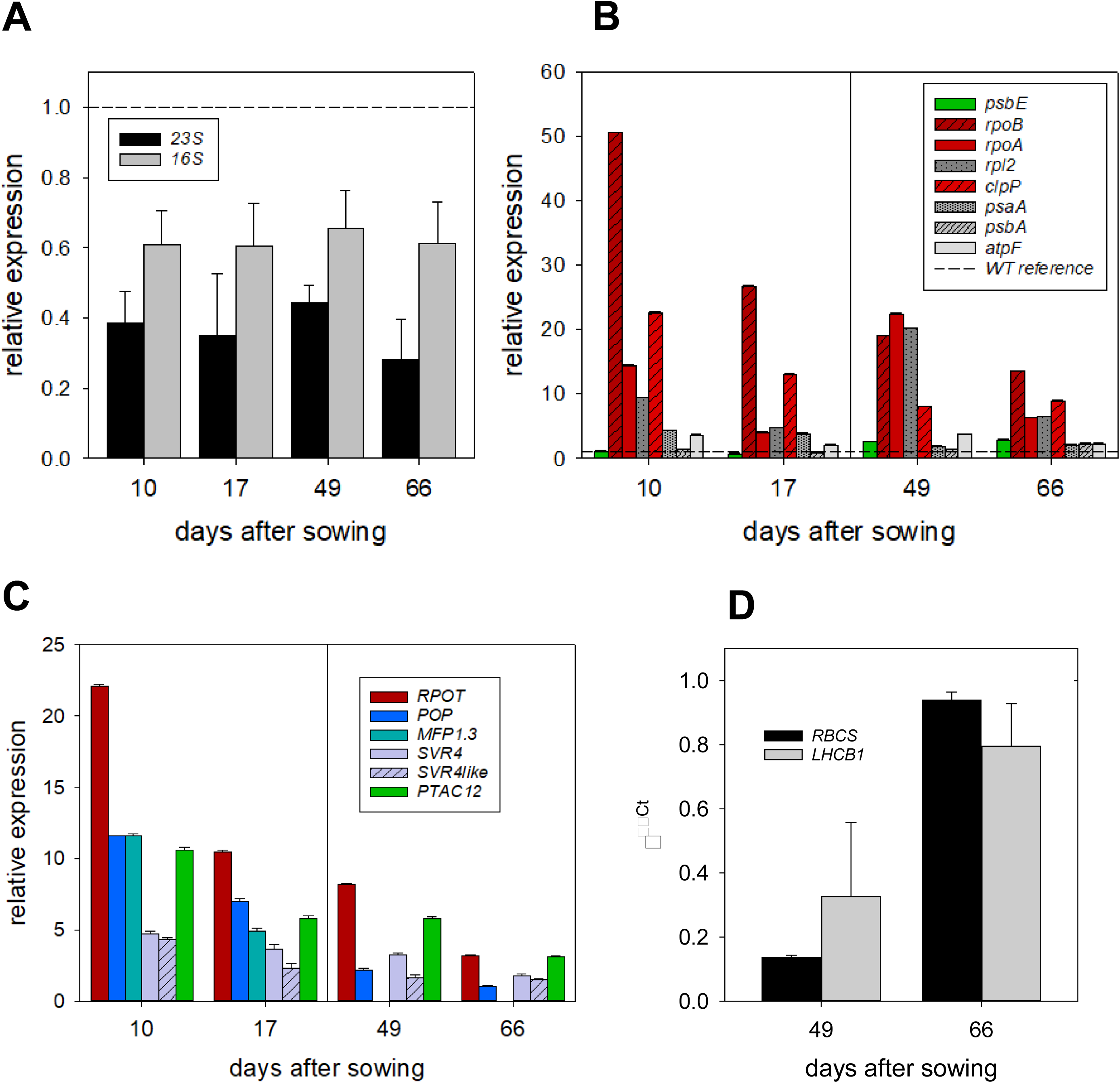
Gene expression in leaves of the wild type and the *why1-3* mutant. **(A)** Relative levels of the large plastid ribosomal RNA species, i.e. 16S and 23S rRNA, **(B)** levels of plastid mRNAs encoding proteins of the photosynthetic apparatus, gene expression machinery and *clpP* encoding the catalytic subunit of the CLP protease complex. Genes transcribed exclusively by PEP are depicted in green, and genes transcribed mainly by NEP are presented in red. Grey columns have been used for genes transcribed by both RNA polymerases. **(C)** Expression levels of nuclear genes encoding the nucleusencoded RNA polymerase of plastids (RPO-T), plant organelle DNA polymerase (POP) and the nucleoid-associated proteins MFP1.3, SVR4, SVR4-like and PTAC12 (PAP5, HEMERA). RNA levels were determined by qRT-PCR using the level of 18S rRNA as standard. The levels of mRNAs in the wild type were set as 1 (interrupted horizontal line).

**Figure 9.**
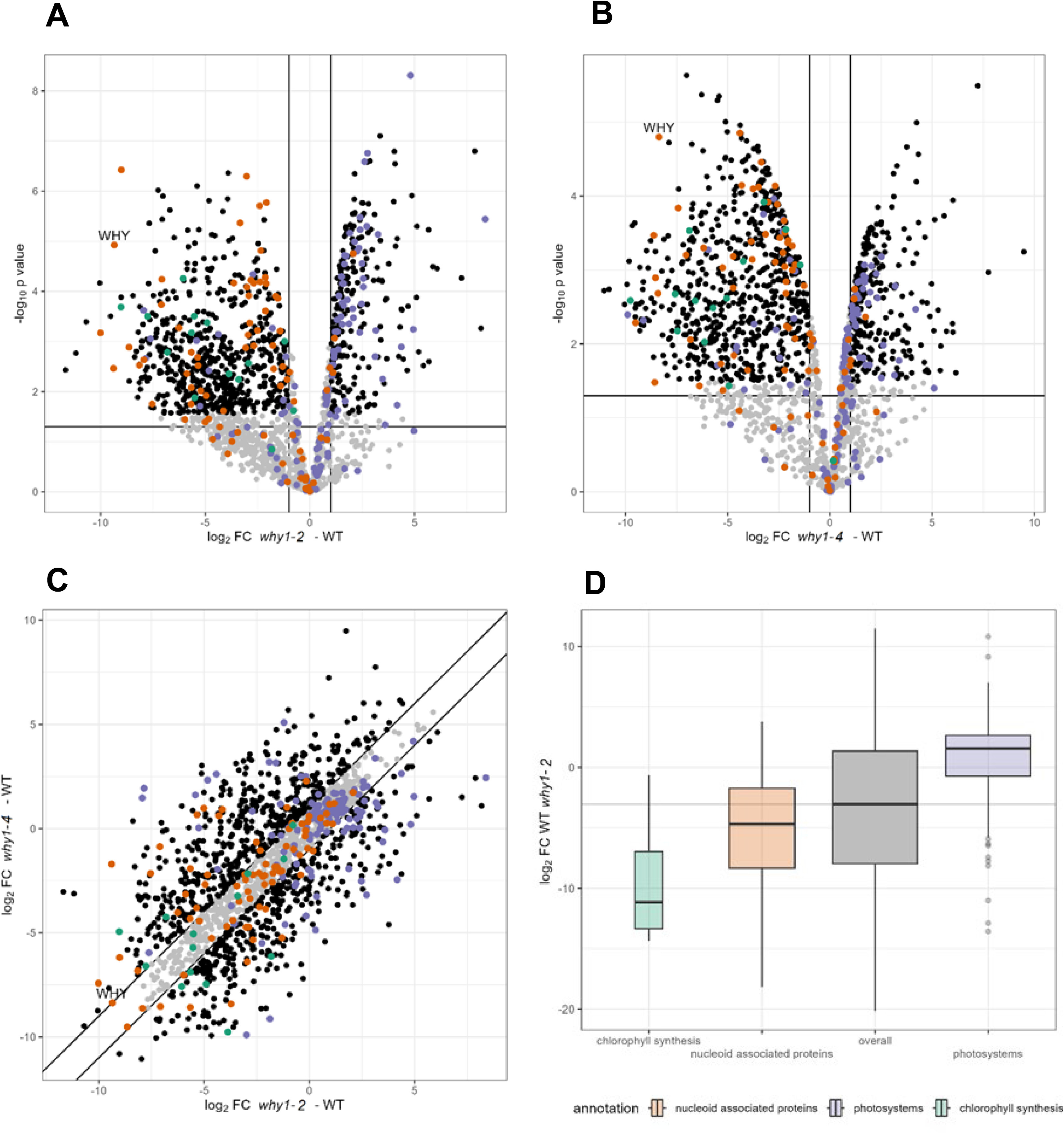
+/- 1) vs -log_10_ p values (horizoProteome analyses of chloroplasts prepared from mature foliage leaves of wild type and the mutants *why1-2* and *why1-4*, respectively. Panels A (*why1-2 - WT*) + B (*why1-4* - WT): Volcano plots correlating LFQ log_2_ fold change (vertical line at zero, horizontal line at 0.05) of limma analysis. Protein groups with adjusted p-value < 0.05 black. Panel C: scatter plot correlating log_2_FC (*why1-3 - WT*) vs log_2_FC (*why1-4* - WT), diagonals and black dots indicating delta FC +/- 1. Panel D: box plots comparing LFQ log_2_ fold change of *why1-11 – WT* overall and per functional assignment. Panels A-D: Photosystem proteins in violet, nucleoid-associated proteins in red and chlorophyll biosynthesis proteins in green.

Both the plastid genes encoding core subunits and the nuclear genes encoding PAP have high gene expression levels. High levels of PAP transcripts are typical for an early stage of chloroplast development preceding the formation of PEP and have also been detected in lethal maize mutants lacking one of the essential nucleus-encoded PAPs such as PTAC12/PAP5 (Kendrick et al., 2022). Although foliage leaves of the barley *why1* mutant are almost as green as wild-type foliage leaves, plastid *rpo* transcripts and nuclear PAP transcripts have elevated levels in these leaves. Possibly, the high levels of the transcripts result from plastid signaling informing about disturbances in transcription and translation (Kendrick et al., 2022).

### Nucleoid compaction in the absence of WHIRLY1

In a previous study with primary foliage leaves of the barley WHIRLY1 knockdown plants (W1-7), chloroplasts had a rather heterogeneous nucleoid population, and small compact nucleoids were observed (Krupinska et al., 2014). In plastids of primary foliage leaves of the *why1* mutant plants, all nucleoids are rather uniform in size but larger than in the wild-type chloroplasts (Figure 6). This indicates that the compaction of nucleoids is reduced in the absence of WHIRLY1. In comparison, chloroplast nucleoids in the mature mutant foliage leaves are almost as small as in wild-type chloroplasts indicating a better compaction. Theoretically, the size of nucleoids stained by the fluorescing dye SYBR®Green could depend on the total amount of ptDNA, which is higher in the *why1* mutant (Figure 6). Since the ptDNA level is, however, higher in foliage leaves than in primary leaves, the level of ptDNA seemingly has no obvious impact on the size of the fluorescence signals.

The higher compaction of nucleoids in foliage leaves indicates that with advancing development the lack of WHIRLY1 is compensated by other proteins. Thereby, the barley *why1* mutant plants can develop chloroplasts that can produce enough assimilates for filling up grains. In order to get insights into the composition of nucleoids and regarding the abundance of nucleoid-associated proteins (NAPs), proteome analyses were performed on isolated chloroplasts of mutant and wild type. By this approach, only a few nucleoid-associated proteins were detected to be upregulated, such as two kinases, the calcium sensor CAS, the MAR attachment region filament-like protein1 (MFP1), and the tetratricopeptide protein 34 (TCP34). The latter two are eukaryotic DNA binding proteins that potentially could have replaced WHIRLY1 in the barley *why1* mutant. Both proteins have their highest abundance early in the chloroplast development (Jeong et al., 2003; Weber et al., 2006). MFP1 had been suggested to have a function in the attachment of nucleoids to both envelope and thylakoid membranes (Jeong et al., 2003) and was suggested to play a role at the interface between nucleoids and the developing thylakoid-membrane system facilitating membrane insertion of photosynthesis-associated proteins and/or linking photosynthesis to gene expression. Recently, MFP1 was shown to interact with a protein involved in initiating starch granules (Seung et al., 2018), suggesting that starch is synthesized in association with nucleoids (Nishimura, 2023). By its structure, MFP1 was suggested to engage in extensive and versatile protein-protein interactions (Sharma et al. 2024). Similar to WHIRLY1, also MFP1 binds to ptDNA without sequence specificity. Both belong to those eukaryotic proteins that have replaced the original prokaryotic nucleoid architects/nucleoid organizing proteins, which have been lost during evolution (Sato, 2001; Kobayashi et al., 2002). The tetratricopeptide-containing chloroplast protein TCP34 (helical repeat proteins) is another eukaryotic protein non-specifically binding to plastid DNA and was proposed to be important for chloroplast development (Weber et al., 2006). Indeed, TCPs were found to transiently participate in almost all processes crucial for thylakoid membrane biogenesis, i.e. stabilization and repair of assembled complexes consisting of proteins, pigments and metal cofactors (Bohne et al., 2016).

Compensation for WHIRLY1 deficiency requires not only an overlap in the functions of WHIRLY1 and the compensatory protein but also similar expression patterns and abundances. WHIRLY1 and MFP1 both have a high abundance in the maize nucleoid, making up 3-5% of the protein mass of nucleoids (Majeran et al., 2012) (relative abundances in Table I: WHIRLY1: 3.7, MFP1: 5.5). However, the abundance of WHIRLY1 on one hand and MFP1 on the other hand in proplastids and chloroplasts show opposing trends. While WHIRLY1 is much more abundant in proplastids (where it has a similar abundance as MFP1) than in chloroplasts (Majeran et al., 2012; Krupinska et al., 2014), abundances of MFP1 and TCP34 increase with chloroplast development (Jeong et al., 2003; Weber et al., 2006; Majeran et al., 2012). Regarding its low abundance in nucleoids (Majeran et al. 2012, Table I: 0.013), TCP34 is unlikely to determine the architecture of nucleoids. It might be rather involved in protein interactions that WHIRLY1 usually mediates.

The severe perturbance in chloroplast development of barley *why1* mutants indicates, however, that the putative compensatory mechanisms cannot fully replace the multiple functions of WHIRLY1 during chloroplast development. Why the compensatory mechanisms are even less efficient in the *why1* mutants of maize and rice mutants remains unknown.

### Nucleoids as assembly platform coordinating chloroplast development

Chloroplast development requires a concerted synthesis and assembly of the photosystems from proteins encoded by plastids and the nucleus as well as numerous co-factors, which all need to be present at the same time (Mullet, 1988). The results of the proteome study on *why1* mutant chloroplasts revealed that WHIRLY1 affects diverse processes required for chloroplast development, such as ribosome formation and chlorophyll biosynthesis and, furthermore, the build-up of ATP synthase, pyruvate dehydrogenase, the CLP protease complex and Rubisco. Considering that the subunits of these complexes are shared by several nucleoid proteomes (Melonek et al., 2016), the nucleoid likely serves as a hub for protein homeostasis and energy production in chloroplasts.

By impacting the compaction of nucleoids, WHIRLY1 strongly impacts all processes taking place at the surface of nucleoids, thereby promoting chloroplast development.

Besides 11 ribosomal proteins detected to be down-regulated in the *why1* mutant chloroplast proteomes, also 16 proteins with different roles in the ribogenesis (Schmid et al., 2023) were less abundant in the *why1* mutant chloroplast proteomes (Supplemental Table 3). These findings are in line with the role of nucleoids as a platform for rRNA processing and ribosome assembly (Bohne, 2014).

The down-regulation of the subunits of pyruvate dehydrogenase might affect lipid biosynthesis in the why1 mutant chloroplasts. The downregulation of the CLP protease might further compromise chloroplast development, which requires efficient repair processes (Richter et al., 2023). Nucleoids contain the chaperon of the CLP protease complex (Melonek et al., 2016). Besides CLPC1, at least 12 CLP protease subunits were less abundant in the mutant chloroplast proteome compared to WT (Table 1). Rubisco is a heterooligomer assembled from plastid-encoded large subunit and the nucleus-encoded small subunit imported into plastids. The assembly of the holo-enzyme depends on chaperonins (Gruber and Feiz, 2018), and its catalytic activity depends on Rubisco activase which belongs to a family of proteins with chaperone-like functions (Portis, 2003). Besides other enzymes involved in the CBB cycle, the Rubisco activating factor RAF1 (Xia et al., 2020) was found to be less abundant in the mutant chloroplast proteome (-5.0, -6.3). Among the proteins associated with carbon fixation, the RbcL showed the strongest decrease (log FC -8.6. -9.5, Table 1) in the mutant chloroplast proteome. Also, the small subunit RBCS (-5.4, -5.0) was found to strongly decrease in abundance compared to WT. Despite the severe reduction in Rubisco, mutant foliage leaves can still perform about 60 % of the CO2 fixation measured for wild-type leaves (Figure 3), confirming that the high abundance of Rubisco is not essential for photosynthesis (Quick et al., 1991; Miller et al., 2000).

Despite the compromised chloroplast development, a functional photosynthetic apparatus is built up in mature foliage leaves of *why1* mutants, and chlorophyll-binding proteins were found even more abundant in the mutant, while the chlorophyll supply was reduced. Although several thylakoid membrane proteins were found to be up-regulated (Figure 8D), there is a dramatic shortage of enzymes required for CO2 fixation, indicating that WHIRLY1 also is required for a balanced composition of the photosynthesis machinery.

Among the proteins down-regulated in the mutant chloroplast proteome are several chaperons which are required for the correct assembly of the multi-subunit complexes down-regulated in the mutant chloroplast proteome. The general chaperone system of chloroplasts consists of ATP-dependent chaperonins and HSP chaperons (Trösch et al. 2015). Both the subunits of the chaperonins (CPN60A, CPN60B, CPN20, CPN10) and the CLPC1 chaperon of the CLP protease showed a decreased abundance in *why1* relative to the wild type. CLPC1 and the chaperonin proteins were identified in immuno-precipitates obtained with maize nucleoids and a specific antibody directed towards ZmWHIRLY1 (Supplementary Table 6 of Majeran et al. 2012). Indeed, of the many subunits of the CLP protease, only the chaperon is found in the catalogue of NAPs indicating that it is tightly attached to the nucleoid, likely by (direct or indirect) interaction with WHIRLY1. Chaperonins were first detected as chaperones mediating Rubisco assembly (Barraclough and Ellis 1980). Later they were found to have roles in assembly at multiple complexes, e.g. ATP synthase, pyruvate dehydrogenase (Blume et al., 2013), and also of ribosomes (Westrich et al., 2021). Among the interactors of the CPN60 machinery, CLPC1 has also been detected, pointing to a close collaboration of the two complexes (Ries et al., 2023). It is proposed that interactions of WHIRLY1 with chaperones mediate the assembly of multi-subunit complexes at the surface of nucleoids (Figure 10).

**Figure 10.**
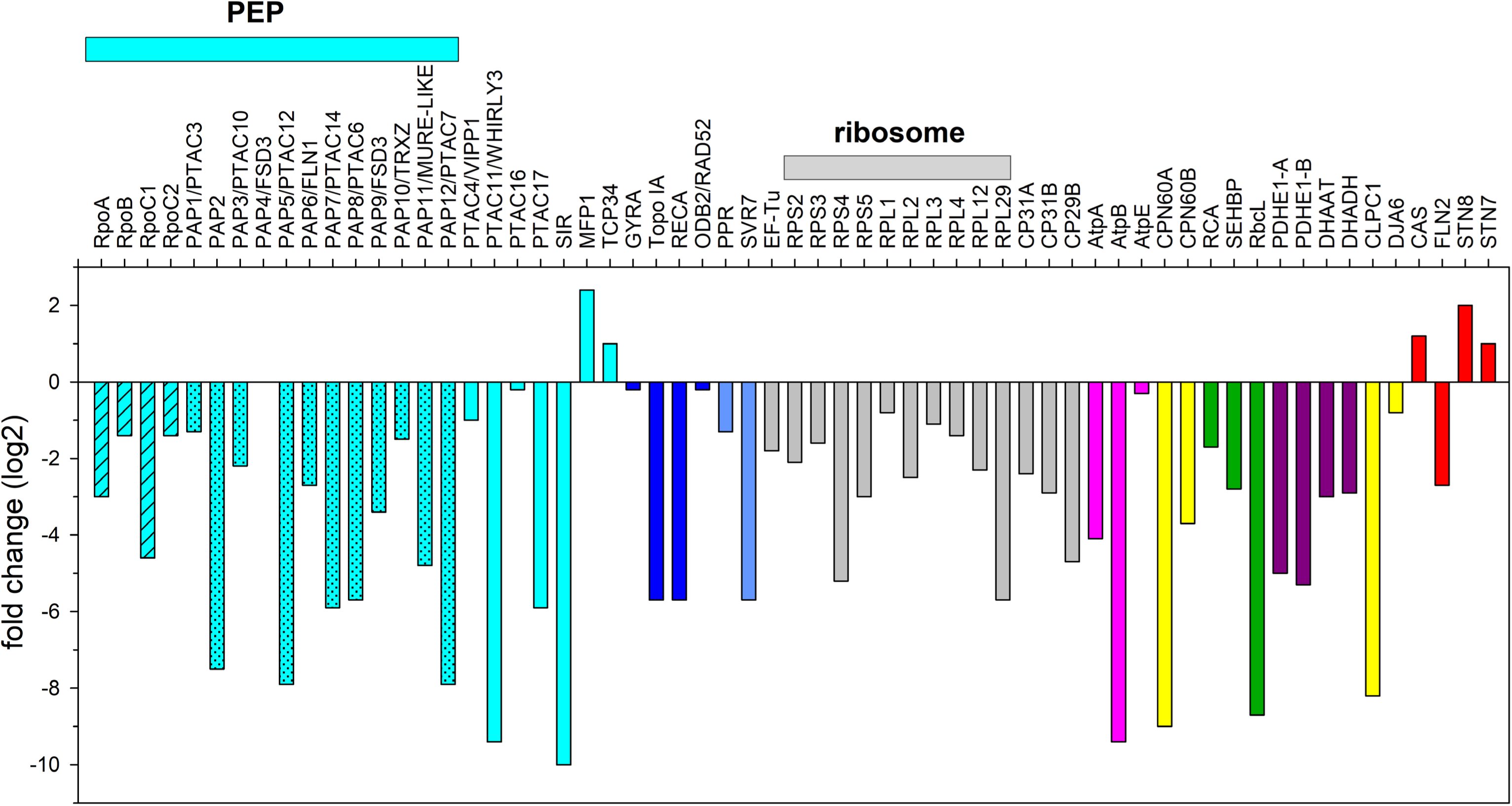
Protein composition of nucleoids as affected by WHIRLY1. Logarithmic fold changes detected for nucleoid associated proteins of the *why1-11* mutant chloroplasts are presented in comparison to abundance of these proteins in wild-type chloroplasts.

**Figure 11.**
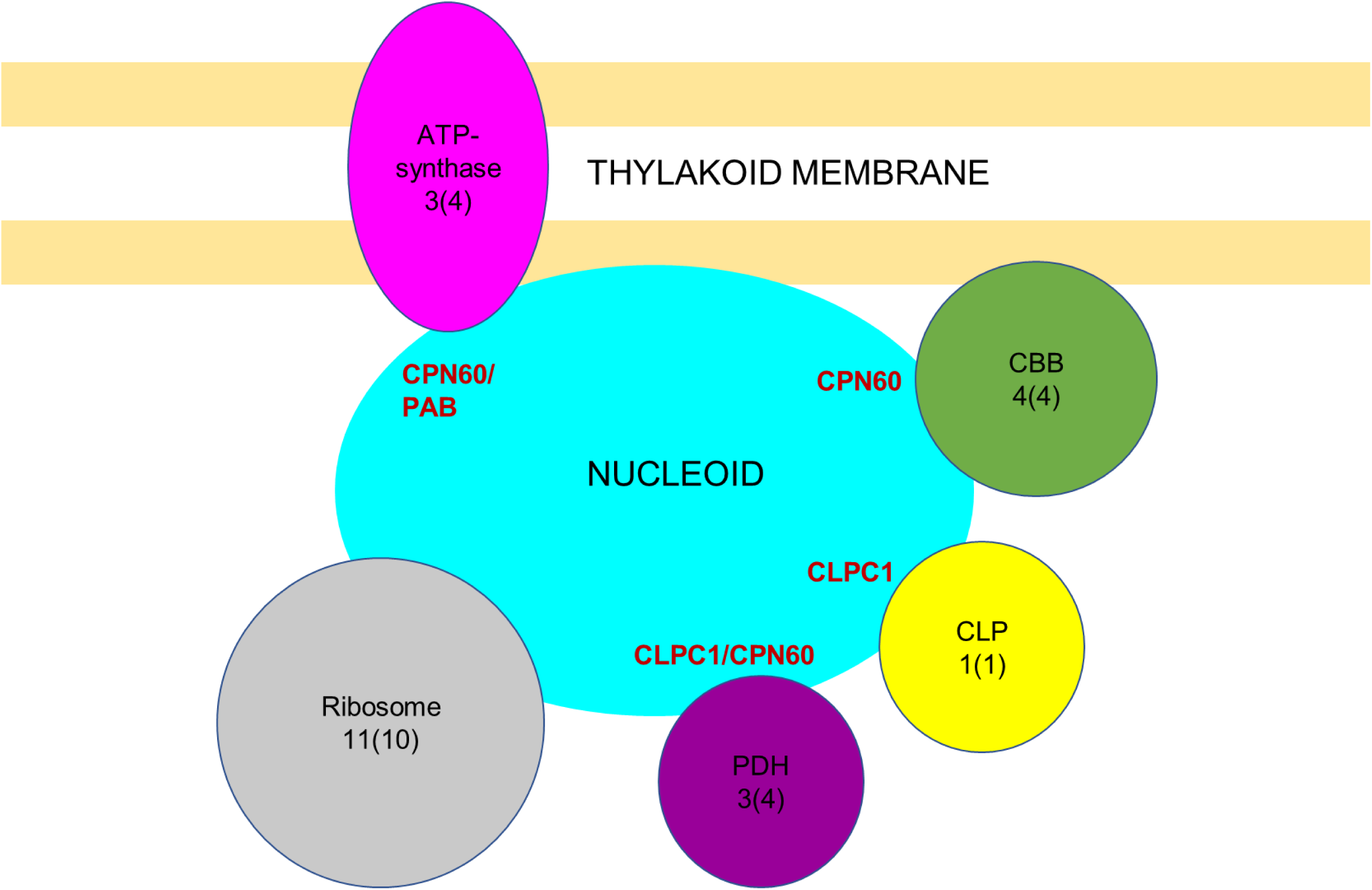
Scheme visualizing the role of WHIRLY1 as an organizer of protein complex assembly in association with nucleoids based on the results presented in Figure 10. The nucleoid is depicted in turquoise in association with the thylakoid membrane (adapted from Krause et al. 2012). Protein complexes containing unexpected nucleoid-associated proteins shared by several nucleoid proteomes (Melonek et al. 2016) and down-regulated in the mutant chloroplast proteome are the ribosomes, pyruvate dehydrogenase (PDH), the CLP protease and the Calvin Bassham Benson cycle (CBB). Chaperones required for assembling the complexes are indicated close to each complex. They are proposed to interact with WHIRLY1 in the periphery of the nucleoid. The number of unexpected nucleoids-associated proteins (UNAPs) downgraded in the mutant chloroplast proteome is indicated for each complex whereby the number in brackets refers to the number of UNAPs shared by several nucleoid proteomes (Melonek et al. 2016), respectively. Colours are assigned to the complexes according to Figure 10.

### No evidence for the involvement of WHIRLY1 in retrograde signaling during chloroplast development

Regarding its dual localization in chloroplasts and nucleus, WHIRLY1 has been proposed as a probable component of retrograde signaling (Grabowski et al., 2008; Bobik and Burch-Smith, 2015). Hence, the barley *why1* mutant is an excellent tool to investigate the role of WHIRLY1 in retrograde signaling evoked by a disturbance of chloroplast development. To address this question, the expression of several nuclear genes known to be controlled by retrograde signaling was analyzed. To begin with, the expression of two photosynthesis-associated nuclear genes (PhANGs), i.e*. LHCB1* and *RBCS*, was analyzed at different developmental stages. It has been reported that the expression of the two genes requires an efficient transcription in the plastids (Rapp and Mullet, 1991). In line with the compromised PEP assembly, the expression of the two genes was reduced (Figure 7).

Furthermore, genes involved in the build-up of the plastidic gene expression machinery, including *RPOT* encoding NEP and the *PAP* genes, are controlled by retrograde signals. Intriguingly, the expression of *PAP* genes was enhanced in the barley *why1* mutant and in chlorotic or albino maize mutants that still possess WHIRLY1 (Kendrick et al., 2022). This indicates that WHIRLY1 is not involved in retrograde signaling targeting these genes. Alice Barkan and co-workers discussed two possibilities for the enhanced expression of PAP genes: on the one hand this could be explained by the lack of a repressive signal originating from photosynthesis, for example, sugars or reducing equivalents, or on the other hand the disruption in chloroplast protein homeostasis (proteostasis) might trigger the increased expression of chloroplast biogenesis genes including *RPOT* and PAP genes (Kendrick et al., 2022). In the barley, *why1* mutant expression of PAP genes is also enhanced in mature foliage leaves capable of photosynthesis. Hence, the second explanation discussed by Alice Barkan and co-workers (see above) seems more plausible. Typically, disruption in proteostasis leads to an up-regulation of genes encoding CLP protease components and chaperones which both are required to restore proteostasis by degradation of aberrant proteins and folding of newly synthesized or imported proteins (Richter et al., 2023). The elevated expression levels of *CLPC1* and *CPN60* in the barley *why1* mutant indicate that the cpUPR is working despite WHIRLY1 is lacking. The reduced abundances of the chaperones in the *why1* chloroplast proteome indicate that proteostasis requires the presence of WHIRLY1. It has been shown that interference of plastid gene expression and activity of the protease CLP, whose enzymatic subunit ClpP is plastome encoded, trigger an unfolded protein response in chloroplasts (cpUPR) (Llamas et al., 2017).

The results of the gene expression analyses indicate that WHIRLY1 is not involved in regulating chloroplast biogenesis genes, PhANGs, or the cpUPR.

### WHIRLY1 might play a role in the control of the plastid DNA level

Leaves of the barley *why1* mutants have an increased level of ptDNA (Figure 6) in line with the enhanced ptDNA levels detected in primary foliage leaves of *HvWHIRLY1* knockdown lines (Krupinska et al., 2014). In the primary foliage leaves of the mutant, the ptDNA level is only slightly more than doubled, which is the level in the W1-7 line having only a minute amount of WHIRLY1. However, the ptDNA content is elevated up to fivefold in the foliage leaves. This result fits the high expression of the gene encoding DNA polymerase 1, which has also been denoted organelle DNA polymerase (POP) (Moriyama et al., 2011; Morley et al., 2019) (Figure 7) (Krupinska et al., 2014). While the expression level of POP in the primary foliage leaves collected at 10 das was more than 10-fold higher than in the wild type, in elder foliage leaves it was only twice as high as in the wild type. Generally, POP expression declined during chloroplast development while the abundance of WHIRLY1 was shown to decrease (Krupinska et al., 2014). Since the abundance of WHIRLY1 in the total protein extract does not inform about its abundance in the nucleus, WHIRLY1 might still participate in the negative regulation of *POP*.

So far, the regulation of DNA copy numbers in plastids has remained largely unknown. In barley, it has been shown that inhibition of transcription has no impact on the plastid DNA copy (Rapp and Mullet, 1991). The copy number is high in actively dividing leaf cells (Nielsen et al., 2010), but is not coupled to the cell cycle. Copy number was decreased with advanced chloroplast development (Boffey et al., 1979; Baumgartner et al., 1989) when the organelle genome faces a higher level of oxidative damage (Mahapatra et al., 2021). It has been shown that oxidatively damaged plastid DNA is degraded by specific glycosylases/endonucleases (Gutman and Niyogi, 2009; Dizdaroglu et al., 2017; Mahapatra et al., 2021). DNA polymerase 1 is not only required for replication, but also for base excision repair (BER). DNA recombination is enhanced in the absence of base excision repair (BER). It is possible that the repair of oxidatively modified bases requires an appropriate structure of nucleoids, which are disorganized in the absence of WHIRLY1. This assumption is in line with the role of WHIRLIES in preventing organelle DNA recombination (Maréchal and Brisson, 2010). An accumulation of *oxidatively* modified DNA might cause a high expression of POP. The molecular mechanism of retrograde signaling by oxidized plastidial DNA leading to a DNA damage response in the nucleus is not known (Mahapatra et al., 2021).

### Conclusions

Due to its dual localization in chloroplasts and nucleus, WHIRLY1 has been suggested as a mediator of retrograde signalling during chloroplast development (biogenic control) (Chan et al., 2016). Obviously, chloroplast development is compromized in the *why1* mutant. Nevertheless, several pathways of retrograde signaling (biogenic control) resulting from disturbances in chloroplast development were shown to be active in the barley *why1* mutant. Therefore, the role of WHIRLY1 in chloroplast development is likely not due to its second localization in the nucleus, where it is known to influence the expression of many genes (Comadira et al., 2015; Krupinska et al., 2022; Taylor et al., 2023). So far, its role in retrograde signaling seems to be specific to stress situations, as shown by investigations with barley plants over-expressing WHIRLY1 (Frank et al., 2023; Manh et al., 2023).

Considering the slow greening of mutant leaves, chloroplast development can, in principle, occur in the absence of WHIRLY1, indicating that WHIRLY1 is not essential for chloroplast development. Rather, WHIRLY1 provides guidance for a concerted workflow of numerous processes required for chloroplast development. Its presence accelerates the process and prevents disturbances of proteostasis by oxidative stress.

## MATERIAL AND METHODS

### Site-directed mutagenesis using Cas9 endonuclease

Twenty nucleotide (nt) target sequences present immediately adjacent to a Protospacer Adjacent Motif (PAM) were selected using DESKGEN^TM^ CRISPR Libraries and verified using the RNAfold web server (http://rna.tbi.univie.ac.at/cgi-bin/RNAWebSuite/RNAfold.cgi) according to Kumlehn et al. (2018).

Oligonucleotides (Supplemental Table X, online) with complementary target sequences and overhangs for cloning into BsaI restriction sites were annealed and inserted between the *OsU3* (RNA polymerase III) promoter and the downstream gRNA scaffold present in the monocot-compatible intermediate vector pSH91 (Budhagatapalli et al., 2016) generating vectors pGH387 and 388. Next, the fragment containing the expression cassettes of gRNA and Cas9 was introduced into the binary vector p6i-d35S-TE9 (DNA-Cloning-Service) using the SfiI restriction sites, generating vectors pGH457 and 458. The vectors constructed for the two target motifs were then used in *Agrobacterium*-mediated barley cv ‘Golden Promise’ co-transformation to generate primary mutant plants. The presence/absence of T-DNA in the regenerated plantlets and mutation detection were achieved by PCR using custom-designed T-DNA–specific and *HvWHIRLY1*-specific primers, respectively.

### Plant material and growth conditions

Barley grains were sown on soil (Einheitserde ED73, Tantau, Ütersen, Germany) and transferred to a climate chamber where the seedlings were grown in a standard daily light/dark cycle (16:8) as described (Krupinska *et al*., 2019).

### Determination of chlorophyll content

To determine the chlorophyll content, one cm long leaf segments excised from the area between 1.5 and 3 cm below the leaf tip were immediately frozen in liquid nitrogen and kept at −80°C. Pigments were extracted, and HPLC analysis was performed as described (Saeid-Nia *et al*., 2022). For non-invasive estimation of the chlorophyll content, a Dualex Scientific instrument (Force A, Paris, France) was used.

### Characterization of photosynthesis

CO2 fixation rates (A) and FV/FM were measured employing a portable Gas Exchange Fluorescence System (GSF-3000, Heinz Walz GmbH, Effeltrich, Germany). FV/FM was determined after keeping the plants for 30-120 min at very low irradiance (PFD = 5 µmol m^-2^ s^-1^) and a further 5 min in darkness in the measuring cuvette. This treatment was sufficient to reach FV/FM values above 0.80 in WT leaves. To determine the carboxylation efficiency and maximal assimilation rate (Amax), attached leaves were first allowed to reach a stable CO2 fixation rate at a PFD of 200 µmol m^-2^ s^-1^ and a CO2 concentration of 400 ppm. Then PFD was increased to 1200 µmol m^-2^ s^-1^, and after a stable assimilation rate was reached, CO2 concentration was raised to 2000 ppm to determine Amax. Subsequently, CO2 concentration was stepwise decreased to 500, 250, 180, 150, 120, 90 and 60 ppm to determine the initial slope of the function of A in dependence on internal CO2 concentration (ci), which is defined as the carboxylation efficiency (CE). A linear regression through the data at the 4 lowest ci was calculated to receive CE. Further conditions during the measurements were 20 °C and 65 % relative humidity.

### Staining and microscopy of nucleoids

Leaf cross-sections were produced by hand or a hand microtome from the primary foliage leaf of plants grown for 7 days. Sections were fixed by 4% (w/v) paraformaldehyde in phosphate-buffered saline (PBS) overnight at 4°C. After washing with PBS containing 0.12 %(w/v) Glycine, the sections were stained with SYBR®Green (1:5000, S7563 Invitrogen^TM^) for 45 min in darkness at room temperature. After washing with 1x PBS for 15 min, the sections were transferred onto a slide, capped with PBS/glycerol (v/v: 1:1), and a coverslip.

Imaging was done at Leica SP5 confocal microscope system with an HCX PL APO CS 63.0 x 1.2 W objective. Excitation was done by an argon laser line 488 (5% power). Emission was detected between 510-570 nm (HV750) and 690-760 nm (HV480). A minimum of five images out of different regions of the specimen were taken from each sample. Image analysis, coloring, and composition were done by ImageJ 1.53q.

### Transmission electron microscopy

Samples were taken from the mid of primary foliage leaves or foliage leaves at a position of two cm below the tip, thereby avoiding the midvein. Resin embedding was performed as described in Saeid Nia et al. (2022). Sections were inspected in a CM10 (Philips) transmission electron microscope, equipped with a MegaView CCD camera (EMSIS), or in a Tecnai G2 Spirat BioTwin transmission electron microscope, equipped with an Eagle 4k × 4k CCD camera (both ThermoFisher Scientific).

### Determination of relative ptDNA levels by quantitative PCR

Total genomic DNA was extracted from barley leaves as described (Fulton et al. 1995). qPCR analyses were performed with a QuantiFast SYBR Green PCR kit (Qiagen, Hilden, Germany) according to the manufacturer’s protocol using specific primers for *petD* and *psbA* as in a previous work (Krupinska et al., 2014). Each reaction was repeated at least three times. Data were normalized to the 18S rDNA gene. The *RBCS* gene was used as a reference for nuclear DNA content.

### Isolation of RNA and determination of mRNA levels by quantitative RT-PCR

Total RNA was isolated from the primary foliage leaves of seedlings or the upper half of foliage leaves using the peqGOLD-TriFast reagent (Peqlab Biotechnology, Erlangen, Germany) according to the manufacturer’s protocol. cDNA biosynthesis and real-time PCR were performed as described previously (Krupinska *et al*., 2019). Data were normalized to the level of the ADP-ribosylation factor 1 mRNA (Rapacz *et al*., 2012), to cytosolic GAPDH in case of RNA from primary foliage leaves or to 18SrRNA in c ase of RNA from foliage leaves.

### Preparation of chloroplasts

Chloroplasts were isolated from foliage leaves by the procedure of Gruissem et al. (1986) using Percoll steps gradients for purification.

### LC-MS/MS and data analysis

Proteins were purified and digested following the SP3 (https://www.nature.com/articles-/s41596-018-0082) procedure with the following adjustments: DTT and IAA were replaced with (tris(2-carboxyethyl)phosphine (TCEP) and 2-chloracetamid (CAA) for reduction and alkylation. Peptide samples were analyzed using an EASY-nLC1200 (Thermo Fisher) coupled to an Exploris 480 (E480) mass spectrometer (Thermo Fisher). Peptides were separated and sprayed with 25 cm fused silica emitters (75 μm inner diameter, CoAnn Technologies) packed in-house with ReproSil-Pur 120 C18 AQ 1.9 μm (Dr. Maisch). 0.5 μg of peptides per sample were analyzed using a stepped gradient of 0% to 45% solvent B (80% ACN, 0.1 % FA) in 60 min at 250 µL/min or 0% to 55% solvent B in 100 min at 300 µL/min, followed by wash steps. Peptide survey mass spectra were acquired in the Orbitrap analyzer, with a resolution of 120,00. A resolution of 15,000 for MS2 spectra was used. The scanned mass range was 300 to 1750 m/z. The normalized collision energy was set to 25. Peptides with a charge of +1, > +6, or peptides with an unassigned charge state were excluded from fragmentation. The respective raw data is available via jPOST under these credentials for review and will be made available under the identifier JPST002385.

URL https://repository.jpostdb.org/preview/1435013137661e52354c413; Access key 3836

Processing of raw data was performed using the MaxQuant software version 2.1.3.0. MS/MS spectra were assigned to the *Hordeum vulgare* NCBI, Uniprot and Phytozome fasta protein databases. During the search, sequences of 248 common contaminant proteins, as well as decoy sequences, were automatically added. Trypsin specificity was required, and a maximum of two missed cleavages was allowed. Carbamidomethylation of cysteine residues was set as fixed, and oxidation of methionine, deamidation, and protein N-terminal acetylation were set as variable modifications. A false discovery rate of 1% for peptide spectrum matches and proteins was applied. Match between runs, as well as LFQ and iBAQ were enabled.

For statistical analyses of the protein abundances, the MaxQuant protein groups table was imported into R. LFQ intensities were log2 transformed. Missing values were imputed using the impute.QRILC function of the imputeLCMD package. Protein groups were considered for further analysis if they were quantified in more than 2 replicates of the respective genotype or condition. Differential enrichment analysis was performed using limma to compare LFQ intensities of WT and the *why1* mutant lines. For functional enrichment analysis, proteins with the respective difference of logFC as value were analysed via the separate function provided by the stringDB.

## ACKNOWLEDGEMENTS

Susanne Braun and Susann Frank (CAU Kiel) are thanked for isolating the chloroplasts used for proteome analyses. Sibylle Freist (IPK Gatersleben) is acknowledged for technical support for generating barley *why1* mutants. Paulina Heinkow (MSPUB, University of Muenster) is acknowledged for maintaining the mass spectrometers and protein sample preparation.

Klaus Humbeck and Ralf-Bernd Klösgen are thanked for their remarkably generous support of the run-on transcription assays carried out at the Institute of Plant Biology of the University of Halle, Germany. We are grateful to Ulrike Voigt (now employed in the Central Microscopy Facility, CAU) and Tanja Zable for technical support in sample preparation for electron microscopy. In addition, we acknowledge the Central Microscopy, CA, for providing Leica SP5 confocal microscopes and transmission electron microscopes.

## Notes

https://repository.jpostdb.org/preview/1435013137661e52354c413

### Competing Interest Statement

The authors have declared no competing interest.

## REFERENCES

Afsson, A.K.E.G. (1938) Studies on the genetic basis of chlorophyll formation and mechanism of induced mutating. Hereditas 24, 33–93

Akoyunoglou, G., Argyroud.Jh, Michelwo.Mr, and Sironval, C. (1966). Effect of intermittent and continuous light on chlorophyll formation in etiolated plants. Physiologia Plantarum 19, 1101-+.

Baumgartner, B.J., Rapp, J.C., and Mullet, J.E. (1989). Plastid transcription activity and DNA copy number increase early in barley chloroplast development. Plant Physiol. 89, 1011–1018.

Bobik, K., and Burch-Smith, T.M. (2015). Chloroplast signaling within, between and beyond cells. Frontiers in Plant Science 6.

Bohne, A.V. (2014). The nucleoid as a site of rRNA processing and ribosome assembly. Frontiers in Plant Science 5.

Börner, T., Aleynikova, A.Y., Zubo, Y.O., and Kusnetsov, V.V. (2015). Chloroplast RNA polymerases: Role in chloroplast biogenesis. Biochimica Biophysica Acta 1847, 761–769.

Budhagatapalli, N., Schedel, S., Gurushidze, M., Pencs, S., Hiekel, S., Rutten, T., Kusch, S., Morbitzer, R., Lahaye, T., Panstruga, R., Kumlehn, J., and Hensel, G. (2016). A simple test for the cleavage activity of customized endonucleases in plants. Plant Methods 12.

Cappadocia, L., Parent, J.S., Zampini, E., Lepage, E., Sygusch, J., and Brisson, N. (2012). A conserved lysine residue of plant Whirly proteins is necessary for higher order protein assembly and protection against DNA damage. Nucleic Acids Research 40, 258–269.

Chan, K.X., Phua, S.Y., Crisp, P., McQuinn, R., and Pogson, B.J. (2016). Learning the languages of the chloroplast: retrograde signaling and beyond. In Annual Review of Plant Biology, Vol 67, S.S. Merchant, ed, pp. 25-+.

Colombo, N., Emanuel, C., Lainez, V., Maldonado, S., Prina, A.R., and Börner, T. (2008). The barley plastome mutant CL2 affects expression of nuclear and chloroplast housekeeping genes in a cell-age dependent manner. Molecular Genetics Genomics 279, 403–414.

De Santis-Maciossek, G., Kofer, W., Bock, A., Schoch, S., Maier, R.M., Wanner, G., Rudiger, W., Koop, H.U., and Herrmann, R.G. (1999). Targeted disruption of the plastid RNA polymerase genes rpoA, B and C1: molecular biology, biochemistry and ultrastructure. Plant Journal 18, 477–489.

de Souza, A., Wang, J.Z., and Dehesh, K. (2017). Retrograde Signals: Integrators of Interorganellar Communication and orchestrators of plant development. In Annual Review of Plant Biology, Vol 68, S.S. Merchant, ed, pp. 85–108.

Demarsy, E., Buhr, F., Lambert, E., and Lerbs-Mache, S. (2012). Characterization of the plastid-specific germination and seedling establishment transcriptional programme. Journal of Experimental Botany 63, 925–939.

De Santis-Maciossek, G., Kofer, W., Bock, A., Schoch, S., Maier, R.M., Wanner, G., Rüdiger, W., Koop, H.U., and Herrmann, R.G. (1999). Targeted disruption of the plastid RNA polymerase genes rpoA, B and C1: molecular biology, biochemistry and ultrastructure. Plant Journal 18, 477–489.

Dillon, S.C., and Dorman, C.J. (2010). Bacterial nucleoid-associated proteins, nucleoid structure and gene expression. Nature Reviews Microbiology 8, 185–195.

Falk, J., Schmidt, A., and Krupinska, K. (1993). Characterization of plastid DNA-transcription in ribosome deficient plastids of heat-bleached barley leaves. Journal of Plant Physiology 141, 176–181.

Fulton, T.M., Chunwongse, J., and Tanksley, S.D. (1995). Microprep protocol for extraction of DNA from tomato and other herbaceous plants. Plant Molecular Biology Reporter 13, 207–209.

Gawronski, P., Burdiak, P., Scharff, L.B., Mielecki, J., Górecka, M., Zaborowska, M., Leister, D., Waszczak, C., and Karpinski, S. (2021). CIA2 and CIA2-LIKE are required for optimal photosynthesis and stress responses in *Arabidopsis thaliana*. Plant Journal 105, 619–638.

Grabowski, E., Miao, Y., Mulisch, M., and Krupinska, K. (2008). Single-stranded DNA binding protein Whirly1 in barley leaves is located in plastids and the nucleus of the same cell. Plant Physiology 147, 1800–1804.

Graciet, E., Gans, P., Wedel, N., Lebreton, S., Camadro, J.M., and Gontero, B. (2003). The small protein CP12: A protein linker for supramolecular complex assembly. Biochemistry 42, 8163–8170.

Jeong, S.Y., Rose, A., and Meier, I. (2003). MFP1 is a thylakoid-associated, nucleoid-binding protein with a coiled-coil structure. Nucleic Acids Research 31, 5175–5185

Kobayashi, K., Narise, T., Sonoike, K., Hashimoto, H., Sato, N., Kondo, M., Nishimura, M., Sato, M., Toyooka, K., Sugimoto, K., Wada, H., Masuda, T., and Ohta, H. (2013). Role of galactolipid biosynthesis in coordinated development of photosynthetic complexes and thylakoid membranes during chloroplast biogenesis in Arabidopsis. Plant Journal 73, 250–261.

Kobayashi, Y., Takusagawa, M., Harada, N., Fukao, Y., Yamaoka, S., Kohchi, T., Hori, K., Ohta, H., Shikanai, T., and Nishimura, Y. (2016). Eukaryotic components remodeled chloroplast nucleoid organization during the green plant evolution. Genome Biology and Evolution 8, 1–16.

Krupinska, K., Desel, C., Frank, S., and Hensel, G. (2022). WHIRLIES are multifunctional DNA-binding proteins with impact on plant development and stress resistance. Frontiers in Plant Science 13.

Krupinska, K., Melonek, J., and Krause, K. (2013). New insights into plastid nucleoid structure and functionality. Planta 237, 653–664.

Krupinska, K., Oetke, S., Desel, C., Mulisch, M., Schäfer, A., Hollmann, J., Kumlehn, J., and Hensel, G. (2014). WHIRLY1 is a major organizer of chloroplast nucleoids. Frontiers in Plant Science 5.

Krupinska, K., Blanco, N.E., Oetke, S., and Zottini, M. (2020). Genome communication in plants mediated by organelle-nucleus-located proteins. Philosophical Transactions of the Royal Society B-Biological Sciences 375.

Krupinska, K., Braun, S., Nia, M.S., Schäfer, A., Hensel, G., and Bilger, W. (2019). The nucleoid-associated protein WHIRLY1 is required for the coordinate assembly of plastid and nucleus-encoded proteins during chloroplast development. Planta 249, 1337–1347.

Kumlehn, J., Pietralla, J., Hensel, G., Pacher, M., and Puchta, H. (2018). The CRISPR/Cas revolution continues: From efficient gene editing for crop breeding to plant synthetic biology. Journal of Integrative Plant Biology 60, 1127–1153.

Li, M.J., Hensel, G., Melzer, M., Junker, A., Tschiersch, H., Ruwe, H., Arend, D., Kumlehn, J., Börner, T., and Stein, N. (2021). Mutation of the ALBOSTRIANS Ohnologous Gene HvCMF3 Impairs Chloroplast Development and Thylakoid Architecture in Barley. Frontiers in Plant Science 12.

Liebers, M., Cozzi, C., Uecker, F., Chambon, L., Blanvillain, R., and Pfannschmidt, T. (2022). Biogenic signals from plastids and their role in chloroplast development. Journal of Experimental Botany 73, 7105–7125.

Liere, K., Weihe, A., and Borner, T. (2011). The transcription machineries of plant mitochondria and chloroplasts: Composition, function, and regulation. J. Plant Physiol. 168, 1345–1360.

Luijsterburg, M.S., Noom, M.C., Wuite, G.J.L., and Dame, R.T. (2006). The architectural role of nucleoid-associated proteins in the organization of bacterial chromatin: A molecular perspective. Journal of Structural Biology 156, 262–272.

Majeran, W., Friso, G., Asakura, Y., Qu, X., Huang, M.S., Ponnala, L., Watkins, K.P., Barkan, A., and van Wijk, K.J. (2012a). Nucleoid-Enriched Proteomes in Developing Plastids and Chloroplasts from Maize Leaves: A New Conceptual Framework for Nucleoid Functions. Plant Physiology 158, 156–189.

Meier, I., Phelan, T., Gruissem, W., Spiker, S., and Schneider, D. (1996). MFP1, a novel plant filament-like protein with affinity for matrix attachment region DNA. Plant Cell 8, 2105–2115.

Melonek, J., Oetke, S., and Krupinska, K. (2016). Multifunctionality of plastid nucleoids as revealed by proteome analyses. Biochim. Biophys. Acta-Proteins and Proteomics 1864, 1016–1038.

Melonek, J., Mulisch, M., Schmitz-Linneweber, C., Grabowski, E., Hensel, G., and Krupinska, K. (2010). Whirly1 in chloroplasts associates with intron containing RNAs and rarely co-localizes with nucleoids. Planta 232, 471–481.

Mullet JE (1993). Dynamic regulation of chloroplast transcription. Plant Physiology 103: 309–313

Nevarez, P.A., Qiu, Y.J., Inoue, H., Yoo, C.Y., Benfey, P.N., Schnell, D.J., and Chen, M. (2017). Mechanism of dual targeting of the phytochrome signaling component HEMERA/pTAC12 to plastids and the nucleus. Plant Physiology 173, 1953–1966.

Nielsen, N.C., Smillie, R.M., Henningsen, K.W., von Wettstein, D., and French, C.S. (1979). Composition and function of thylakoid membranes from grana-rich and grana-deficient chloroplast mutants of barley. Plant Physiology 63, 174–182.

Oetke, S., Scheidig, A., and Krupinska, K. (2022). WHIRLY1 of barley and maize share a PRAPP motif conferring nucleoid compaction. Plant Cell Physiology 63, 234–247.

Pfalz, J., and Pfannschmidt, T. (2013). Essential nucleoid proteins in early chloroplast development. Trends in Plant Science 18, 186–194.

Pfalz, J., Liere, K., Kandlbinder, A., Dietz, K.-J., and Oelmüller, R. (2006). pTAC2, -6, and -12 are components of the transcriptionally active plastid chromosome that are required for plastid gene expression. Plant Cell 18, 176–197.

Pfannschmidt, T., Nilsson, A., and Allen, J.F. (1999). Photosynthetic control of chloroplast gene expression. Nature 397, 625–628.

Pfannschmidt, T., Terry, M.J., Van Aken, O., and Quiros, P.M. (2020). Retrograde signals from endosymbiotic organelles: a common control principle in eukaryotic cells. Philosophical Transactions of the Royal Society B-Biological Sciences 375.

Pfannschmidt, T., Blanvillain, R., Merendino, L., Courtois, F., Chevalier, F., Liebers, M., Grubler, B., Hommel, E., and Lerbs-Mache, S. (2015). Plastid RNA polymerases: orchestration of enzymes with different evolutionary origins controls chloroplast biogenesis during the plant life cycle. Journal of Experimental Botany 66, 6957–6973.

Pogson, B.J., Woo, N.S., Förster, B., and Small, I.D. (2008). Plastid signalling to the nucleus and beyond. Trends in Plant Science 13, 602–609.

Powikrowska, M., Oetke, S., Jensen, P.E., and Krupinska, K. (2014). Dynamic composition, shaping and organization of plastid nucleoids. Frontiers in Plant Science 5.

Prikryl, J., Watkins, K.P., Friso, G., van Wijk, K.J., and Barkan, A. (2008). A member of the Whirly family is a multifunctional RNA- and DNA-binding protein that is essential for chloroplast biogenesis. Nucleic Acids Research 36, 5152–5165.

Qiu, Z.N., Chen, D.D., Teng, L.H., Guan, P.Y., Yu, G.P., Zhang, P.L., Song, J., Zeng, Q.C., and Zhu, L. (2022). OsWHY1 interacts with OsTRX z and is essential for early chloroplast development in rice. Rice 15.

Rapp, J.C., Baumgartner, B.J., and Mullet, J. (1992). Quantitative analysis of transcription and RNA levels of 15 barley chloroplast genes - transcription rates and messenger-RNA levels vary over 300-fold - predicted messenger-RNA stabilities vary 30-fold. Journal of Biological Chemistry 267, 21404–21411.

Reiter, B., Rosenhammer, L., Marino, G., Geimer, S., Leister, D., and Rühle, T. (2023). CGL160-mediated recruitment of the coupling factor CF 1 is required for efficient thylakoid ATP synthase assembly, photosynthesis, and chloroplast development in Arabidopsis. Plant Cell 35, 488–509.

Rotasperti, L., Sansoni, F., Mizzotti, C., Tadini, L., and Pesaresi, P. (2020). Barley’s second spring as a model organism for chloroplast research. Plants-Basel 9.

Rottet, S., Besagni, C., and Kessler, F. (2015). The role of plastoglobules in thylakoid lipid remodeling during plant development. Biochimica Et Biophysica Acta-Bioenergetics 1847, 889–899.

Saeid Nia, M., Repnik, U., Krupinska, K., and Bilger, W. (2022). The plastid-nucleus localized DNA-binding protein WHIRLY1 is required for acclimation of barley leaves to high light. Planta 25

Sato, N., Hirama, T., Miyajima, K., Sekine, K., Kabeya, Y., and Ehira, S. (2002). Comparative biochemistry of plastid nucleoid proteins - a hypothesis on the discontinuous evolution of plastid genomic machinery. Plant and Cell Physiology 43, S81–S81.

Schmid, L.M., Manavski, N., Chi, W., and Meurer, J. (2023). Chloroplast Ribosome Biogenesis Factors. Plant and Cell Physiology

Taylor, R.E., West, C.E., and Foyer, C.H. (2023). WHIRLY protein functions in plants. Food and Energy Security 12.

Trösch, R., Mühlhaus, T., Schroda, M., and Willmund, F. (2015). ATP-dependent molecular chaperones in plastids - More complex than expected. Biochimica Et Biophysica Acta-Bioenergetics 1847, 872–888.

von Wettstein, D. (1959). Developmental changes in chloroplasts and their genetic control. Symp. Soc. Study Develop. Growth 16, 123–160.

von Wettstein, D. (2001). Discovery of a protein required for photosynthetic membrane assembly. Proceedings of the National Academy of Sciences of the United States of America 98, 3633–3635.

Wang, S.H., and Blumwald, E. (2014). Stress-induced chloroplast degradation in *Arabidopsis* is regulated via a process independent of autophagy and senescence-associated vacuoles. Plant Cell 26, 4875–4888.

Xia, L.Y., Jiang, Y.L., Kong, W.W., Sun, H., Li, W.F., Chen, Y.X., and Zhou, C.Z. (2020). Molecular basis for the assembly of RuBisCO assisted by the chaperone Raf1. Nature Plants 6, 708-+.

Xie, Q.J., Michaeli, S., Peled-Zehavi, N., and Galili, G. (2015). Chloroplast degradation: one organelle, multiple degradation pathways. Trends in Plant Science 20, 264–265.

Zhelyazkova, P., Sharma, C.M., Forstner, K.U., Liere, K., Vogel, J., and Borner, T. (2012). The primary transcriptome of barley chloroplasts: numerous noncoding RNAs and the dominating role of the plastid-encoded RNA polymerase. Plant Cell 24, 123–136.

