## Supplemental Figures for "The major nucleoid-associated protein WHIRLY1 promotes chloroplast development in barley"

| | gRNA1 | *PAM* | Genomic position: |
|---|---|---|---|
| **guide:** | GTAGTCGGCGGAGTGGCGCG | *CGG* | |
| | G**A**AGTCGGCGGAG**A**GGCGCG | *GGG* | CABVVH010000005.1:408214535-408214557 |
| | G**A**AGTCGGCGGAG**A**GGCGCG | *GGG* | CABVVH010000005.1:408321596-408321618 |
| | GTAG**G**CGGCGG**C**GTGGCGCG | *AGG* | CABVVH010000004.1:445851542-445851564 |
| | **gRNA2** | ***PAM*** | **Genomic position:** |
| **guide:** | TCCCTCTCCAGCGGCGGCGG | *CGG* | |
| **off-target:** | TCCCTCTCC**G**GCGGCGGCGG | *CGG* | CABVVH010000006.1:21072245-21072267 |
| | TCCCTCTCCAGCGGC**T**GCGG | *TGA* | CABVVH010000006.1:305667919-305667941 |
| | TCCCTCTCCAGCGGC**T**GCGG | *TGA* | CABVVH010000003.1:249848829-249848851 |
| | T**T**CCTCTCCAGCGGCG**A**CGG | *TGG* | CABVVH010000001.1:168746351-168746373 |
| | TCCCT**T**TCC**T**GCGGCGGCGG | *GGG* | CABVVH010000001.1:432785104-432785126 |
| | TCC**TC**CTCCAGCGGCGGCGG | *CGG* | CABVVH010000001.1:327593303-327593325 |
| | **C**C**A**CTCTCCAGCGGCGGCGG | *CGG* | CABVVH010000004.1:366535973-366535995 |
| | T**T**C**T**TCTCCAGCGGCGGCGG | *CGG* | CABVVH010000005.1:223281033-223281055 |
| | T**G**CCTCTCC**T**GCGGCGGCGG | *CGG* | CABVVH010000002.1:411526964-411526986 |
| | TCC**T**TCTCC**G**GCGGCGGCGG | *CGG* | CABVVH010000001.1:266307189-266307211 |
| | TCCC**A**CTC**T**AGCGGCGGCGG | *CGG* | CABVVH010000002.1:506792562-506792584 |
| | T**A**CCTCTCC**G**GCGGCGGCGG | *TGG* | CABVVH010000008.1:98258377-98258399 |
| | T**A**CCTCTCC**G**GCGGCGGCGG | *TGG* | CABVVH010000008.1:73236038-73236060 |
| | TCC**G**TCTCCA**T**CGGCGGCGG | *GGG* | CABVVH010000007.1:439703643-439703665 |

**Supplementary Figure 1A.** Off-target analysis for gRNAs. Examples with 1 or 2 mismatches (highlighted in bold and red) were shown. Mismatches in the seed region, which are the twelve nucleotides in front of the PAM (underlined), abolish gRNA-Cas9 binding.

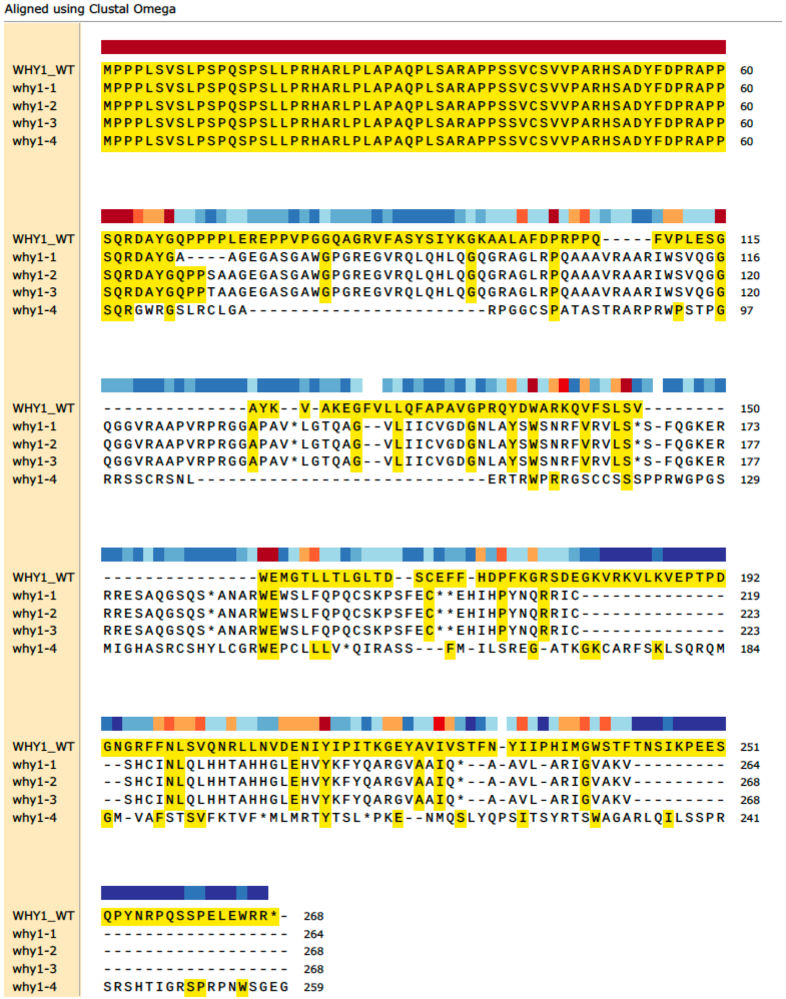

**Supplemental Figure 1B**. A CLUSTALW comparison of the deduced amino acid sequences of WHY1 wildtype and all four mutant plants. The identical amino acids are highlighted in yellow. The colour code on top of the sequence indicates the conservation, with red representing the highest similarity and blue representing the lowest.

**A**

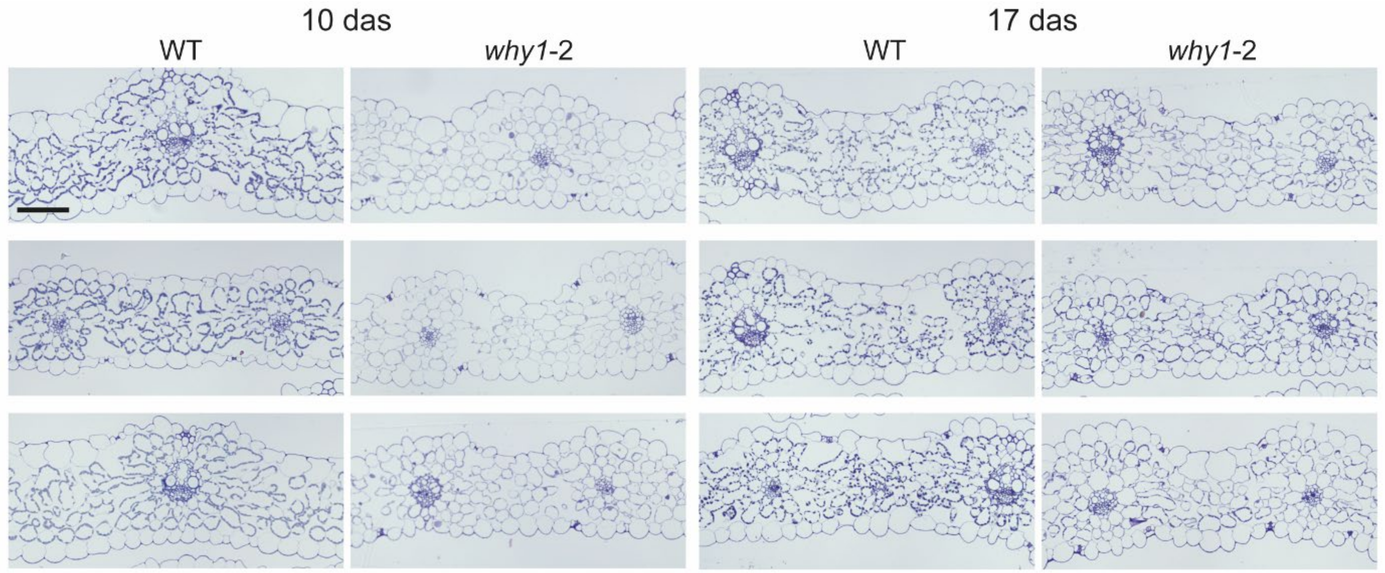

**Supplementary Figure 2A.** Electron microscopy images of semithin **c**ross leaf sections from barley wild type and mutant *why1-11:* **(A)** Primary foliage leaves collected at 10 or 17 days after sowing, respectively. Three leaves were collected for each genotype.
Scalebar, 100 µm.

**B**

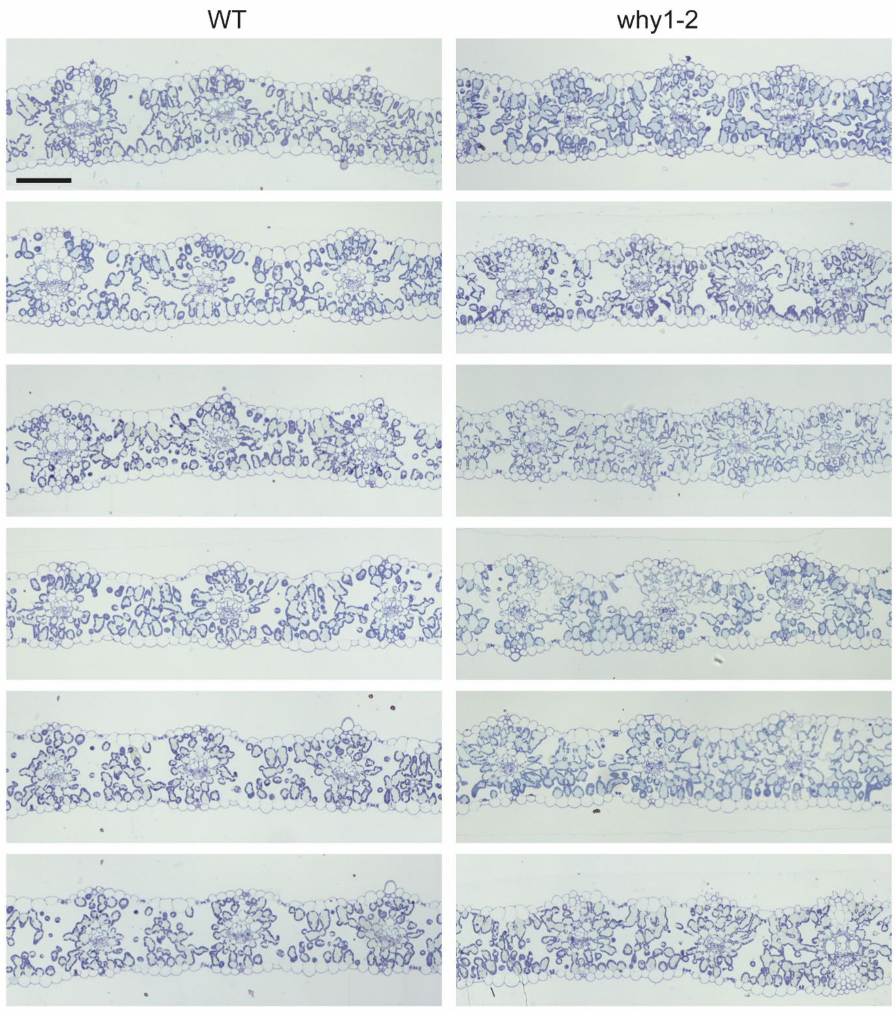

**Supplemental Figure 2B**. Electron microscopy images of semithin cross sections of barley wild type and mutant *why1-2* mature foliage leaves collected three months after sowing. Scalebar, 100 µm.

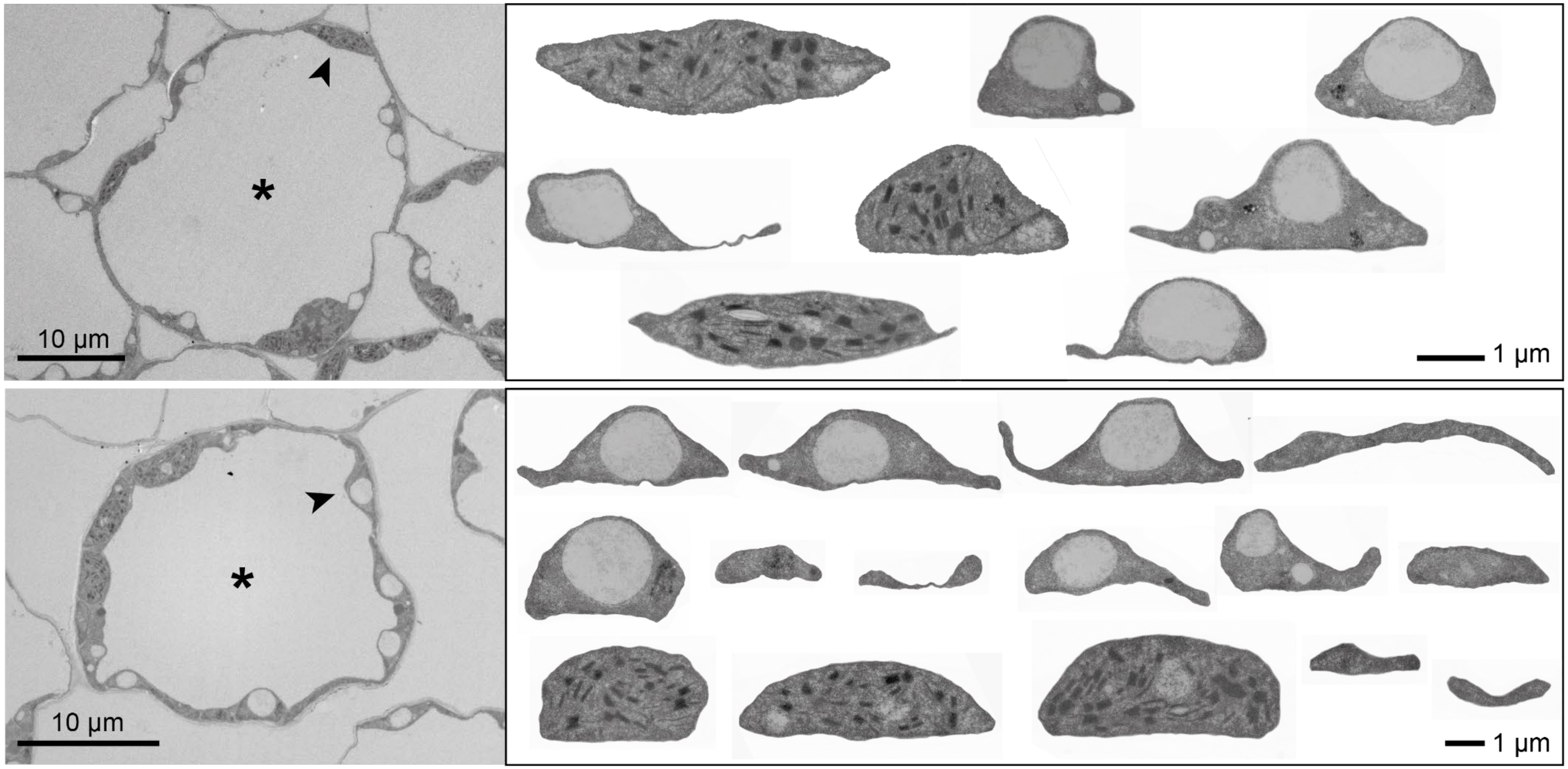

**Supplemental Figure 3.** Electron microscopy images obtained from ultrathin sections of primary foliage leaves of the barley mutant *why1-11.* Images on the left show two exemplaric mesophyll cells with heterogeneous population of plastids, respectively. Starting at the positions indicated by an arrow, the plastids are shown in higher magnification on the right. In the first and third row, respectively, plastids are arranged from left to right, in the middle plastids are arranged from right to left.

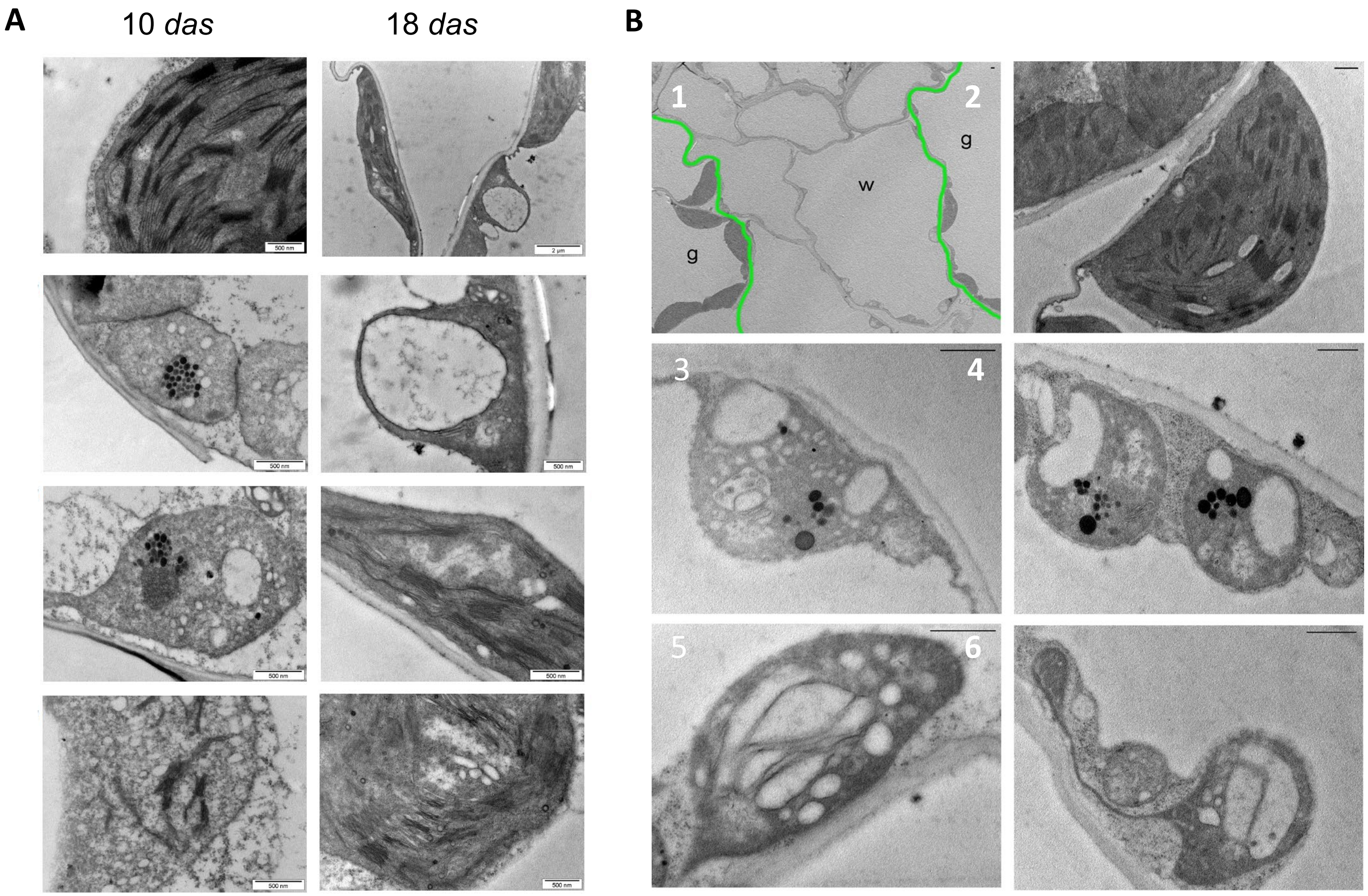

**Supplemental Figure 4**. Ultrastructural analyses of plastids found in barley mutant leaves: **(A)** Plastids observed in primary foliage leaves of the mutant *why1-4* collected at either 10 das or 18 days after sowing (das).
**(B)** Plastids in green/white striped leaves of the barley mutant *albostrians.* (1) Overview showing a white sector in the middle and two green sectors left and right thereof. The borders are indicated by green lines, (2) chloroplast in the green sector, (3-6) different types of plastids in the white sector.
Scale bar in 2-6: 500 nm

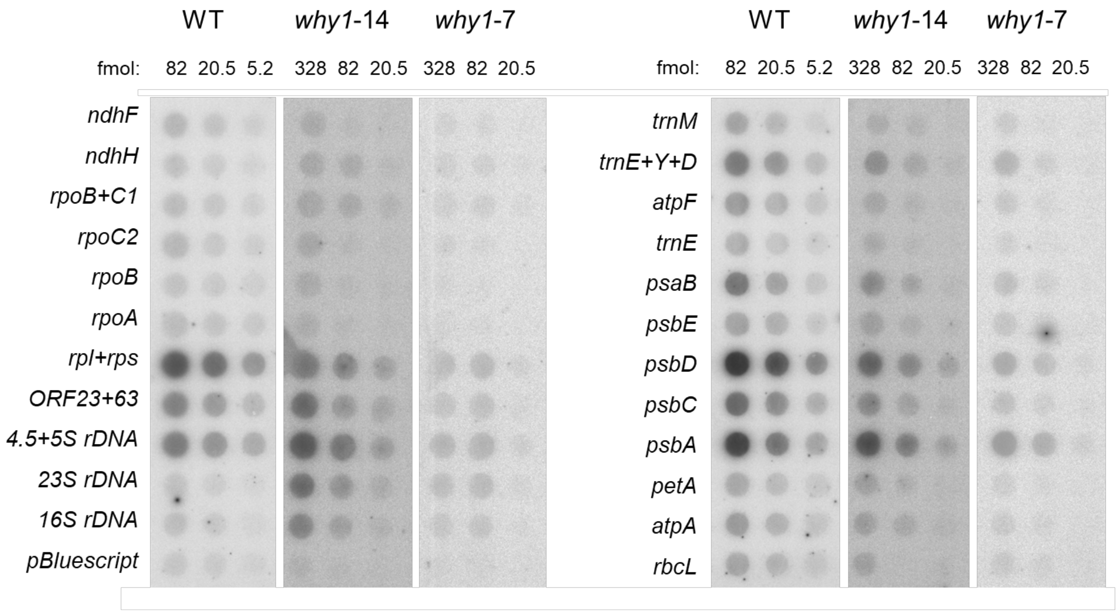

| run on transcripts | WT | *why1*-14 | why*1-7* |
|---|---|---|---|
| *psbE/rbcL* | 1.1 | 2.2 | 1.1 |
| *ndhH/rpoB+C1* | 0.7 | 0.9 | 0.7 |
| *16S rDNA/23S rDNA* | 1.6 | 0.6 | 0.6 |
| rel. transcriptional activity (%) | 100 | 21 | n.d. |

**Supplemental Figure 5.** Run-on transcription assays perfomed with isolated plastids prepared from primary foliage leaves of the wild type (WT) collected 7 days after sowing (*das)* and the *why1-2* mutant collected either 7 (*why1-7*) or 14 *das (why1-14).*
Transcription assays and hybridization of transcripts with specific plastid gene probes were done as desbribed previously (Falk et al. 1993, Melonek et al. 2010). Plasmids with 23 specific plastid genes were dotted on nylon filters in four different concentrations corresponding to 328, 82, 20.5 and 5.2 fmol of each gene specific probe. As a control pBluescript was dotted onto the filter. The DNA dot-blots were hybridized with $^{32}$P-labelled run-on transcripts obtained with wild-type chloroplasts (WT) and mutant plastids (*why1-7, why1-14*).
After hybridization, the filters were exposed to phosphorimager screens and analysed with the Fujifilm FLA-3000 (Fujifilm, Düsseldorf, Germany) using the software packages BASReader (version 3.14) and AIDA (version 3.25; Raytest, Straubenhardt, Germany). For easier visual comparison of hybridization signals, the strong signals obtained by hybridization of wild-type run-on transcripts with 328 fmol probes are not shown, whereas the weak signals obtained by hybridization of mutant run-on transcripts with 5.2 fmol probes are not shown.

The **Table** shows the ratios of relative signal intensities of specific genes. All these calculation were performed with the signals detected with 82 fmol of each gene specific probe. In barley *psbE* and *ndhH* are mainly transcribed by PEP, while the *rpoB/C1* is exclusively transcribed by NEP. *rbcL* and the rRNA gene are transcribed by both RNAPs (Zhelyazkova et al., 2012). The relative transcriptional activities were estimated by adding the signal values of all 23 gene specific probes. This comparison was only possible between the run-on assays performed with the same number of wild-type and mutant plastids (*why1-14*). The number of plastids was not be determined in case of the *why1-7* sample (n.d.).

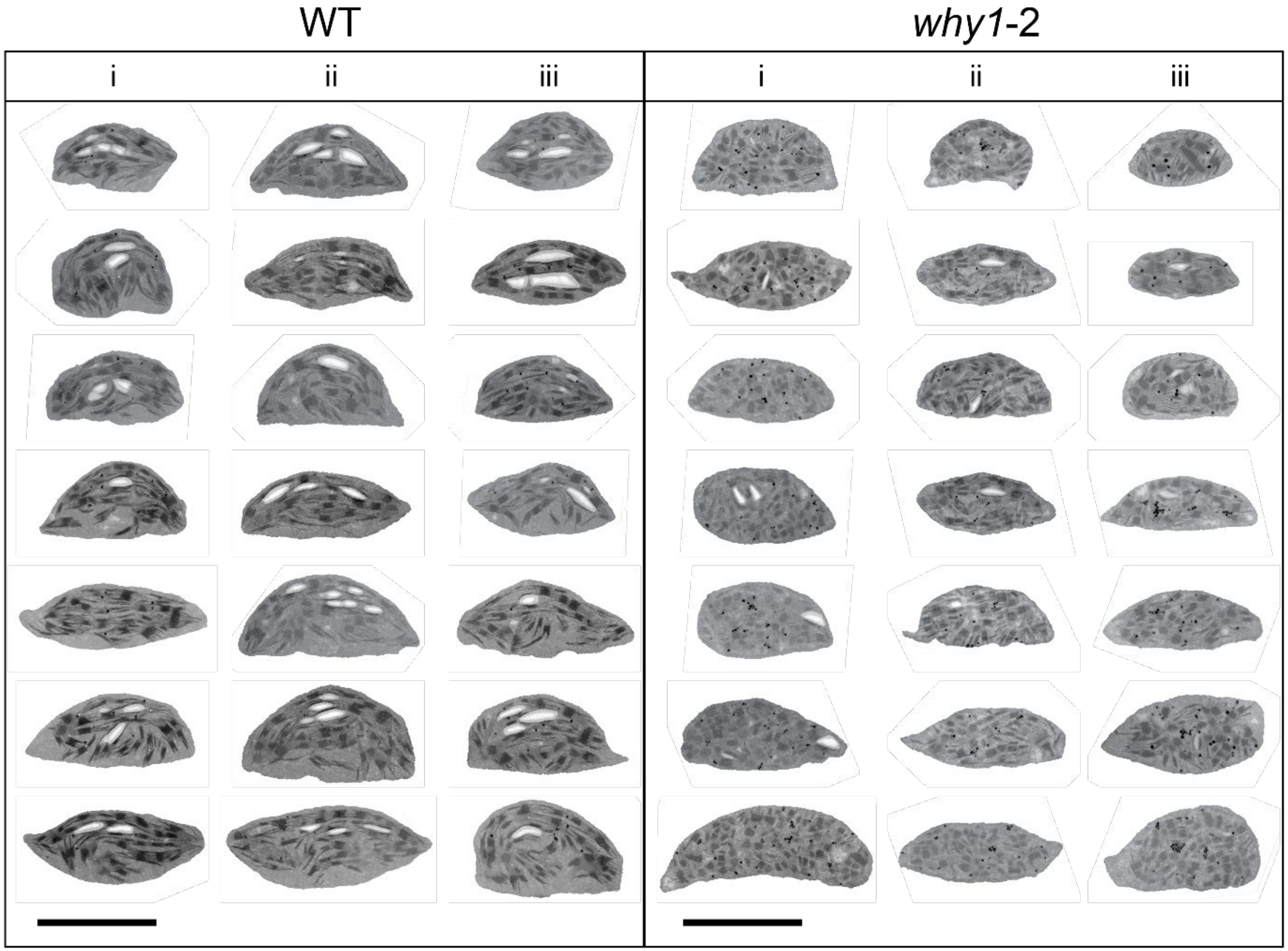

**Supplemental Figure 6.** Comparison of chloroplasts in mesophyll cells of foliage leaves collected from WT and *why1*-2 mutant plants, at 3 months after sowing. Three plants were sampled for each genotype. They were embedded in epoxy resin and chloroplasts were imaged on 80-nm thin sections. Chloroplasts in the mutant leaves seem a little smaller and show less parallel organisation of thylakoids, have more plastoglobules and less starch grains. Scalebar, 5 µm.

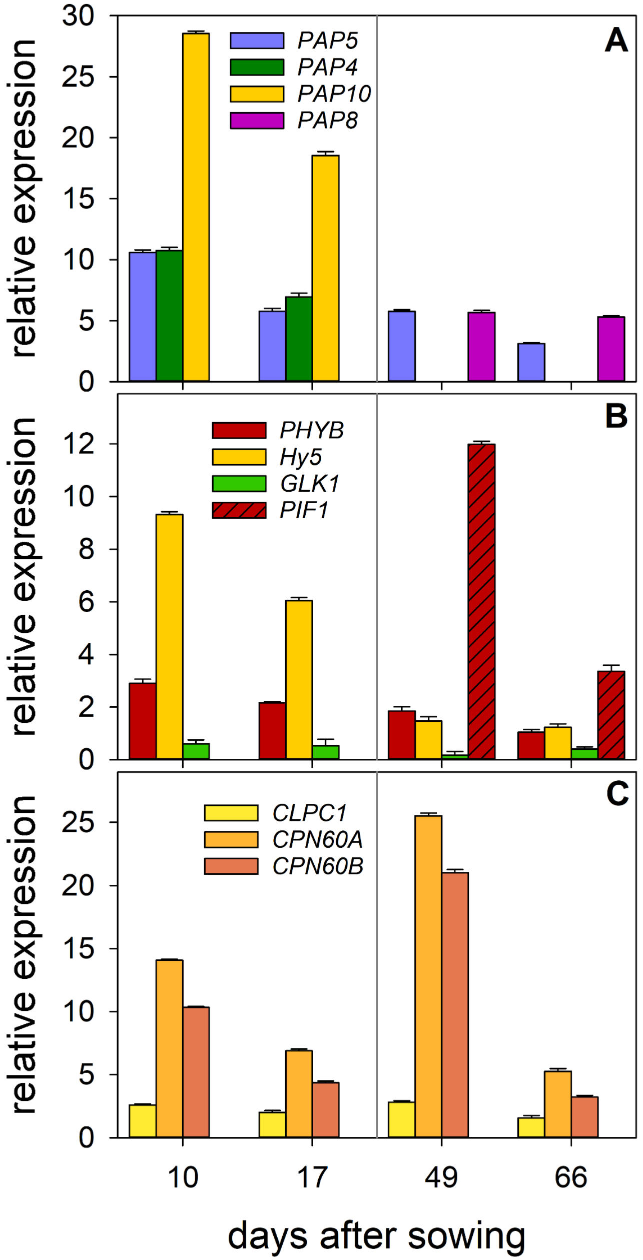

**Supplemental Figure 7.** Expression of nuclear genes in primary foliage leaves of the wild type and the *why1-11* mutant. **(A)** Relative mRNA levels of genes encoding PEP associated proteins PAP5, PAP8 and PAP12, **(B)** relative mRNA levels of genes encoding PHYTOCHROME B (PHYB) and transcription factors HY5, GLK1 and PIF1, respectively, **(C)** mRNA levels of CLPC1 and chaperonin subunits CPN60 A and B. RNA levels were determined by qRT-PCR using the level of *GAPDH* mRNA as standard. The levels of mRNAs in the wild type were set as 1.

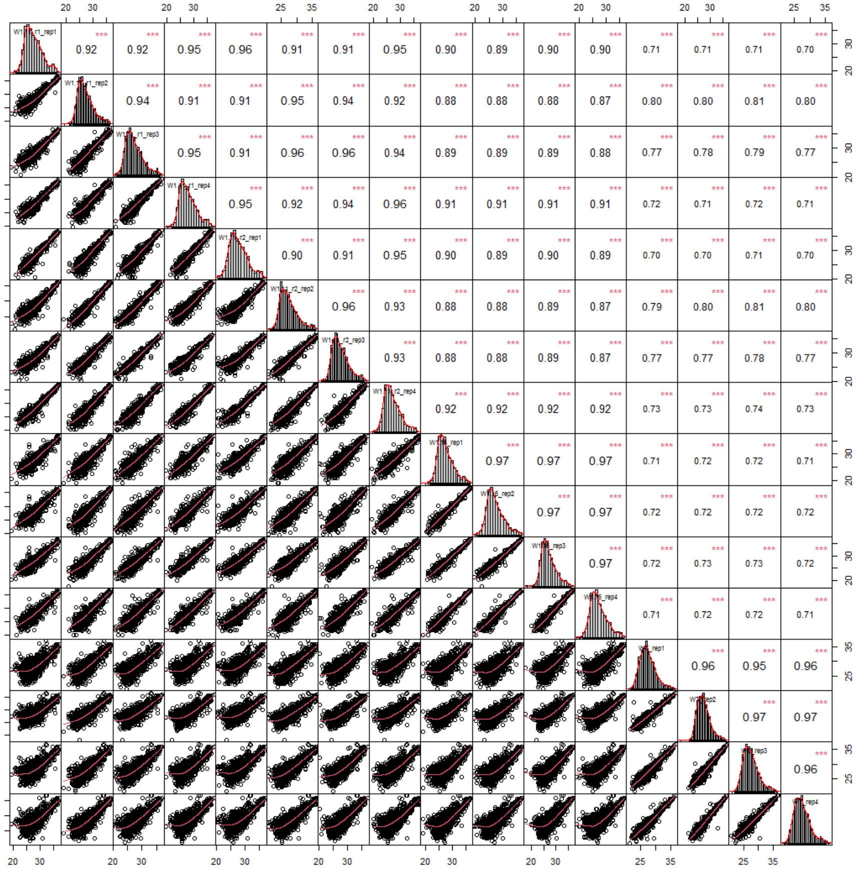

**Supplemental Figure 8.**
Multiscatter plots correlating label free quantification values for all replicates (1-4) and conditions (*why1-2* (2 independent cultivations), *why1-4* and WT) across the dataset.
