## Supplemental Table 1 for "The major nucleoid-associated protein WHIRLY1 promotes chloroplast development in barley"

**Supplemental Table 1.** Catalogue of nucleoid associated proteins (Melonek et a. 2016) and their abundance in the chloroplast proteome of the barley *why1-11* mutant.

n.d. not detected, d. detected but not scored

| Proteome analysis | | | Pfalz et al. 2006 | Melonek et al. 2012 | Phinney and Thelen 2005 | Majeran et al. 2012 | | | Huang et al. 2013 | |
|---|---|---|---|---|---|---|---|---|---|---|
| Number of proteins identified | | *why1*-11 CP proteome logFC | 35 (26 proteins in both mustard and Arabidopsis; 4 only in Arabidopsis and 5 in mustard) | 53 | 179 (35 known or predicted ptNAPs) | 1092 (crude nucleoid fraction) | 888 immuno-precipitated with WHIRLY1 | 157 (core nucleoid proteins) | 1028 (crude nucleoid fraction | 212 (core nucleoid proteins) |
| | **Identified ptNAPs** | | | | | | | | | |
| **Gene name** | **AGI** | | | | | | | | | |
| | ***Subunits of the plastid-encoded RNA polymerase (PEP)*** | | | | | | | | | |
| RpoA | ATCG00740 | -3.0 | + | (+) | + | + | + | + | + | + |
| RpoB | ATCG00190 | -1.4 | + | - | + | + | + | + | + | + |
| RpoC1 | ATCG00180 | -4.6 | + | + | - | + | + | + | + | + |
| RpoC2 | ATCG00170 | -1.4 | + | + | - | + | - | + | + | + |
| Sigma 2 | AT1G08540 | | - | - | - | + | + | - | - | - |
| | ***PEP-associated proteins (PAPs) and additional TAC-components*** | | | | | | | | | |
| MFP1 | AT3G16000 | +2.4 | - | - | - | + | + | - | + | - |
| PEND | AT3G52170 | | - | - | + | - | - | - | - | - |
| PTAC-1 (WHIRLY1) | AT1G14410 | WHIRLY3 | + | - | - | + | + | - | + | + |
| PTAC-2 (PAP2) | AT1G74850 | -7.5 | + | + | + | + | + | + | + | + |
| PTAC-3 (PAP1) | AT3G04260 | -1.3 | + | + | - | + | + | + | + | + |
| PTAC-4 (Vipp1) | AT1G65260 | -1.0 | + | - | - | + | + | + | + | + |
| pTAC-5 (peptidoglycan | AT4G13670 | n.d. | + | - | - | + | + | - | + | + |

| | | | | | | | | | | |
|---|---|---|---|---|---|---|---|---|---|---|
| binding domain-containing protein, DnaJ) | | | | | | | | | | |
| pTAC-6 (PAP8) | AT1G21600 | -5.7 | + | - | + | + | + | + | + | + |
| pTAC-7 (PAP12) | AT5G24314 | -7.9 | + | - | - | + | + | - | + | + |
| pTAC-8 | AT2G46820 | n.d. | + | - | - | + | - | - | - | - |
| pTAC-9 (Organellar single-stranded OSB2) | AT4G20010 | n.d. | + | - | - | - | - | - | - | - |
| pTAC-10 (PAP3) PDE312 | AT3G48500 | -2.2 | + | - | - | + | + | + | + | + |
| pTAC-11 (WHIRLY3) | AT2G02740 | -9.3 | + | - | - | + | - | - | + | + |
| pTAC-12 (PAP5) | AT2G34640 | d. | + | - | - | + | + | + | + | + |
| pTAC-13 | AT3G09210 | d. | + | - | - | + | + | + | + | + |
| pTAC-14 (PAP7) | AT4G20130 | -5.9 | + | - | + | + | + | + | + | + |
| pTAC-15 | AT5G54180 | n.d. | + | - | - | - | - | - | + | + |
| pTAC-16 | AT3G46780 | -0.2 | + | - | - | + | + | - | + | - |
| pTAC-17 | AT1G80480 | -5.9 | + | - | - | + | + | - | + | - |
| pTAC-18 | AT2G32180 | n.d. | + | - | - | + | - | - | + | - |
| MurE-like (PAP11) | AT1G63680 | -4.8 | + | - | - | + | + | + | + | + |
| SiR (DCP68) | AT5G04590 | -10.0 | - | - | + | - | - | - | + | - |
| SWIB-3 | AT4G34290 | n.d. | - | - | - | + | (+) | - | - | - |
| SWIB-4 | AT3G03590 | n.d. | - | - | - | - | (+) | - | + | - |
| SWIB-6 | AT2G35605 | n.d. | - | + | - | - | - | - | - | - |
| TCP34 | AT3G26580 | +1.0 | - | - | - | + | - | - | - | - |
| | ***DNA replication, repair and recombination*** | | | | | | | | | |
| DNA polymerase gamma I (PolIA) | AT1G50840 | n.d. | - | - | - | + | + | - | - | - |
| DNA polymerase 2 (PolIB) | AT3G20540 | n.d. | + | - | - | - | - | - | - | - |

| | | | | | | | | | | |
|---|---|---|---|---|---|---|---|---|---|---|
| Topoisomerase DNA gyrase A (GYRA) | AT3G10690 | -0.2 | + | - | + | + | + | + | + | + |
| Topoisomerase DNA gyrase B (GYRB1) | AT3G10270 | n.d. | + | - | + | - | + | - | + | + |
| Topoisomerase IA | AT4G31210 | -5.7 | - | - | - | + | + | + | + | + |
| DNA exonuleaseI 5'-3'EXO | AT3G52050 | d. | - | - | + | + | + | - | - | - |
| Recombinase (RECA) | AT1G79050 | -5.7 | - | - | + | + | + | - | + | + |
| Organellar-DNA-binding-protein2 (ODB2, RAD52) Recombination mediator | AT5G47870 | -0.2 | - | - | - | + | - | - | + | + |
| | ***Posttranscriptional processes*** | | | | | | | | | |
| At5g46580 PPR | At5g46580 | -1.3 | - | + | - | + | + | + | + | + |
| SVR7 (Suppressor of variegation 7) | AT4G16390 | -5.7 | - | - | - | + | - | + | + | - |
| RAP (RNA binding domain abundant in apicomplexans) | AT2G31890 | n.d. | - | - | - | + | + | + | + | - |
| Zm-mTERF4 | AT4G02990 | n.d. | - | - | - | + | + | + | + | + |
| ZmTERF9 | AT5G55580 | n.d. | - | - | - | + | + | - | - | - |
| | ***Translation*** | | | | | | | | | |
| EF-Tu | AT4G20360 | -1.8 | + | + | + | + | + | - | + | - |
| 30S ribosome S2 | ATCG00160 | -2.1 | - | - | + | + | + | - | + | + |
| 30S ribosome S3 | ATCG00800 | -1.6 | + | - | - | + | + | + | + | + |
| 30S ribosome S4 | ATCG00380 | -5.2 | - | - | + | + | + | + | + | + |
| 30S ribosome S5 | AT2G33800 | -3.0 | - | + | - | + | + | - | + | + |
| 50S ribosome L1 | AT3G63490 | -0.8 | - | - | + | + | + | + | + | + |
| 50S ribosome L2 | ATCG01310 | -2.5 | - | - | + | + | + | + | + | + |

| | | | | | | | | | | |
|---|---|---|---|---|---|---|---|---|---|---|
| 50S ribosome L3 | AT2G43030 | -1.1 | - | - | + | + | + | + | + | + |
| 50S ribosome L4 | AT1G07320 | -1.4 | - | - | + | + | + | + | + | + |
| 50S ribosome L12 | AT3G27850 | -2.3 | + | - | - | + | + | - | + | - |
| 50S ribosome L15 | AT3G25920 | ? | - | - | + | + | + | + | + | + |
| 50S ribosome L29 | AT5G65220 | -5.7 | + | - | | + | + | + | + | - |
| CP31A | AT4G24770 | -2.4 | - | - | + | + | + | - | + | - |
| CP31B/28 kDa ribonucleoprotein spinach | AT5G50250/ CAA41023 | -2.9 | - | - | + | + | + | - | + | - |
| CP29B/24 kDa ribonucleoprotein spinach, RRM | AT2G37220/ AAA79045 | -4.7 | - | - | - | + | + | - | + | - |
| | ***ATP synthesis*** | | | | | | | | | |
| alpha subunit (AtpA) | ATCG00120 | -4.1 | + | + | + | + | - | - | + | - |
| beta subunit (AtpB) | ATCG00480 | -9.4 | + | + | + | + | + | - | + | - |
| gamma subunit (ATPC) | AT4G04640 | n.d. | - | + | + | + | + | - | + | - |
| epsilon subunit (AtpE) | ATCG00470 | -0.3 | - | + | - | + | - | - | + | - |
| | ***Acetyl-CoA carboxylase*** | | | | | | | | | |
| alpha subunit (ACC1) | AT1G36160 | n.d. | - | - | + | + | + | - | - | - |
| transferase beta subunit (AccD) | ATCG00500 | n.d. | - | - | + | - | - | - | + | + |
| biotin carboxylase precursor (CAC2) | AT5G35360 | n.d. | - | - | + | - | - | - | + | - |
| biotin carboxyl carrier protein (CAC1) | AT5G16390 | n.d. | - | - | + | - | - | - | - | - |
| | ***Calvin cycle*** | | | | | | | | | |
| Rubisco binding protein alpha subunit (CPN60A) | AT2G28000 | -9.0 | - | - | + | + | + | - | + | - |

| | | | | | | | | | | |
|---|---|---|---|---|---|---|---|---|---|---|
| Rubisco binding protein beta subunit (CPN60B) | AT3G13470 | -3.7 | - | - | + | + | + | - | + | - |
| Rubisco activase (RCA) | AT2G39730 | -1.7 | + | + | + | + | + | - | + | - |
| seduheptulose-1,7-biphosphatase | AT3G55800 | -2.8 | - | + | + | + | - | - | + | - |
| Rubisco large subunit (LSU) | ATCG00490 | -8.7 | - | - | + | + | + | - | + | - |
| | ***Pyruvatdehydrogenase*** | | | | | | | | | |
| E1 alpha subunit | AT1G01090 | -5.0 | - | - | + | + | + | - | + | - |
| E1 beta subunit | AT1G30120 | -5.3 | - | - | + | + | + | - | + | + |
| dihydrolipoamide S-acetyltransferase | AT1G34430 | -3.0 | + | - | + | + | + | - | + | + |
| dihydrolipoamide dehydrogenase | AT3G25860 | -2.9 | - | - | + | + | + | + | + | - |
| ***Redox proteins*** | | | | | | | | | | |
| FeSOD-1/FSD2 (PAP9) | AT5G51100 | -3.4 | + | - | - | + | + | + | + | + |
| FeSOD-3/FSD3 (PAP4) | AT5G23310 | d. | + | - | - | + | + | + | + | + |
| Thioredoxin Z (TrxZ/PAP10) | AT3G06730 | -1.5 | + | - | - | + | + | + | + | + |
| | ***Protein homeostasis/regulation*** | | | | | | | | | |
| CLP-C1 | AT5G50920 | -8.2 | - | + | + | + | + | - | + | - |
| HSP40 | AT1G09260 | n.d. | - | + | - | - | - | - | - | - |
| HSP70 | AT4G24280 | n.d. | + | + | + | + | + | - | + | - |
| Dja6 (DNA-J PROTEIN A6) | AT2G22360 | -0.8 | - | - | - | + | + | + | + | + |
| | ***Kinases*** | | | | | | | | | |
| pfkB-type carbohydrate kinase (FLN1/PAP6) | AT3G54090 | -2.7 | + | - | - | + | + | + | + | + |

| | | | | | | | | | | |
|---|---|---|---|---|---|---|---|---|---|---|
| PFK-B-type carbohydrate kinase (FLN2) | AT1G69200 | -2.7 | + | - | + | + | + | + | + | + |
| STN8 (State transition 8) | AT5G01920 | +2.0 | - | - | - | + | - | - | + | + |
| STN7 (State transition 7) | AT1G68830 | +1.0 | - | - | - | + | + | + | + | + |
| | ***Diverse processes*** | | | | | | | | | |
| Branched chain amino acid transferase | AT3G05190 | ? | - | + | - | - | + | - | - | - |
| CAS (Calcium sensing receptor) | AT5G23060 | +1.2 | - | - | - | + | + | - | + | + |
| Polygalacturonase | AT3G07820 | n.d. | - | + | - | - | - | - | - | - |
| Mg-chelatase subunit | At4G18480 | n.d. | - | + | + | - | + | - | + | - |
