## Supplemental Table 3 for "The major nucleoid-associated protein WHIRLY1 promotes chloroplast development in barley"

**Supplemental Table 3**. Factors involved in chloroplast ribosome biogenesis (ribogenesis), found in Table 1 of Schmid et al. (2023) (includes 42 proteins), RP=ribosomal protein

| **protein** | **Arabidopsis identifier** | **$log_2$ FC in *why1*-11 and *why1*-16 chloroplasts** | **activity, domain** | **proposed function in ribogenesis** |
|---|---|---|---|---|
| PNPase | AT3G03710 | -2.6, -3.1 | exoribonuclease | maturation of 23S rRNA |
| RNE | AT2G04270 | -5.2, -4.4 | endonuclease | production of the 23S-4.5S rRNA precursor |
| RNR1 | AT5G02250 | -4.0, -2.5 | ribonuclease | trimming of rRNAs |
| RNJ | AT5G63420 | -2.8, -3.8 | ribonuclease | rRNA termini control |
| CSP41a/b | AT3G63140 | -1.6, -2.5 | endoribonuclease | processing of hidden breaks in 16 and 23SrRNAs |
| YbeY | AT2g25870 | -4.0, -3.9 | endoribonuclease | maturation of rRNA ends |
| ISE2 | AT1G7007 | -3.0, -0.1 | RNA helicase | RNA editing and splicing of mRNAs for RPs |
| RHON1 | AT1G06190 | -7.2, -4.7 | RNA-helicase | rRNA processing, guides RNE |
| WSL6(Era-1) | AT5G66470 | -1.8, -1.3 | GTPase | 30S assembly |
| SDP | AT1G12800 | -7.2, -6.8 | RNA chaperon | processing of 23S-4.5S |
| SRRP1 | AT3G23700 | -2.5, -1.1 | RNA chaperon | splicing of *trnL* intron, cleavage of 5S rRNA |
| RBF1 | AT4G34730 | -0.9, -0.3 | RBF A domain | maturation of 16S rRNA ends |
| SOT1 | AT5G46580 | -1.3, -5.2 | PPR-SMR protein | protects 23S-4.5SrRNA precursor |
| DCL | AT1G45230 | -4.9, -3.7 | DUF3223 domain | generates 4.5 SrRNA |
| CRASS | AT5G14910 | -4.8, -6.5 | HMA domain | associates with 16SrRNA and RPs |
| SVR1 | AT2G39140 | -0.9, -3.1 | Pseudouridin synthase | pseudouridylation of rRNAs |
