## Supplemental Table 4 for "The major nucleoid-associated protein WHIRLY1 promotes chloroplast development in barley"

Supplemental Table S4

| Primer | 5' - 3' for | 5' - 3' rev | bp |
|---|---|---|---|
| ***ADP*** | cgtgacgctgtgttgcttgt | ccgcattcatcgcattagg | 61bp |
| ***plastid*** | | | |
| ***atpF*** | caaagggcaatgaatcaggt | agcacgaatcgtacggaaat | 114bp |
| ***rpoA*** | gtatccatgcctgttcgaaat | cgcttccttaggggttaaact | 120bp |
| ***rpoB*** | ggtcgacgaaatatgaatcg | ccaatcaaatgatccgtagc | 101bp |
| ***psbE*** | cgggttggttatttgtcagt | agaatcaaaacggtcggtta | 119bp |
| ***psbA*** | ctgcttggcctgtagtagga | cgcgaccttgactatcaact | 111bp |
| ***psaA*** | cttttcctaatcgcaggtca | aagacccttatggccttgtc | 114bp |
| ***clpP*** | tatccgaagacatggaaagg | ggatcagtatcgcagtgctt | 112bp |
| ***petD*** | GGGCGTTCTCTTAATGGTTT | AATGGGTAGTGTTGCTCCAA | **161bp** |
| ***psbA*** | CAGAAAAGCTTCCTTGACCA | CAATGGTGGTCCTTATGAGC | **187bp** |
| ***rpl2*** | ggtgctgtagcgaaactcat | tacccacttgtccgactgtt | 115bp |
| ***clpP*** | tatccgaagacatggaaagg | ggatcagtatcgcagtgctt | 112bp |
| ***18S*** | caggtccagacatagcaaggattgacag | taagaagctagctgcggagggatgg | 174 bp |
| ***16S*** | gaataagcatcggctaactctg | acttgaaaagccacctacagac | 105bp |
| ***23S*** | ttaaactgcctgctgaatcc | tactacgggaatcgcttttg | 101bp |
| ***nucleus*** | | | |
| ***HY5*** | AAGAAGAATTCGGAGCTGGAAG | TCTGTGCTATTGACCCTCACTT | 123bp |
| ***CLPC1*** | TTGTGTCGTCATGGACTGGTA | TTGCTTTCACAGCCTCATCTT | 123bp |
| ***GLK1*** | GGACATGGAATTACCAGAAGG | CTTGTTCTTGTCGTTGTCGTG | 100bp |
| ***CPN60A*** | TGTCCGCTATTCATTGTAGCC | TCGCCGCAACGTTAATAATAC | 94bp |
| ***CPN60B*** | GAAAATGACCACCGAGTACGA | CCTAATGGCTTCCTCCAGAAC | 100bp |
| ***LHCA1*** | CCGGAGAAGAAGAAGTACCC | CGTTCTTGATCTCCTTGAGC | 100bp |
| ***LHCA3*** | GCAGTACTTCCTGGGTCTCG | CATCTCCTTCTCGCTCTTGG | 112bp |
| ***LHCB1*** | TATCGGCTCTCTTTGGTGAG | AGGTAGAGCACACGATCAGC | 109bp |
| ***LHCB4*** | ACGGCAACACCCTCAACTAC | GGTGGAGCTTGTCATCGAAT | 138bp |
| ***PAP4*** | AAGGATTTCGGCTCCTTTGT | GTATGAACCACCGCGAGTTT | 125bp |
| ***PAP5*** | GCCCGAGTCACTAGAGGATG | TCCTTTGGTTTTTCGTCCTG | 117bp |
| ***PAP8*** | GAGTGGACGAAATCGAGACA | CAAGCTTCCTGCATAAACGA | 116bp |
| ***PAP10*** | CGATCGAAATGGTCAGGAAC | CGGACTGCGCCTAAATAAAC | 121bp |
| ***PHYB*** | TTCCTTGCAGCTCATATTGC | CATCTATTCCCCGCAATTCT | 119bp |
| ***PIF1*** | TTATGCAGCAGGACGTTGAG | CTCCGAGAGGTTGTGCACTT | 110bp |
| ***POP*** | TCCACCATCCAGTACATCCA | CAGTTGGCCATGGAGATGTC | 113bp |
| ***RBCS*** | CTACCACCGTCGCACCCTTCC | TGATCCTTCCGCCATTGCTGAC | 104bp |
| *RPOT* | AAATATGGTGGAGGCATTGA | AACAGTTCCCATCAACAGGA | 1112bp |
| ***MFP1*** | CGAGTTGAACAAGGAATTGGA | ACCTCAGTTCGGGAATCACTT | 123bp |
| ***SVR4*** | CGGACTACTTCGACAAGCAT | CCACTCCAAGCAGTTGATCT | 113bp |
| ***SVR4 like*** | CAGGGATGTCAAGACAAAGG | CCATCTCATTGTCCTGTGGT | 107bp |
| ***PTAC12*** | GCCCGAGTCACTAGAGGATG | TCCTTTGGTTTTTCGTCCTG | 117bp |

| additional information |
|---|

| qPCR nDNA: ptDNA |
|---|

| CPN60A_chr.2 |
|---|
| CPN60B_LIN1 chr.7 |

| PAP4_FSD3 |
|---|
| PAP5_pTAC12 HEMERA |

| PAP10_TRXZ |
|---|

| rpoT (ex9) |
|---|
| MFP1.3 |

| PAP5_pTAC12_HEMERA |
|---|
